# Denoising single-cell ATAC-seq data with Shannon entropy

**DOI:** 10.64898/2026.10.20.760589

**Authors:** Shiyi Yang, Kevin An, Juliana J. Lee, Jase Gehring, Sina Booeshaghi, Haiyan Huang, Aaron Streets

## Abstract

Single-cell Assay for Transposase Accessible Chromatin with sequencing (scATAC-seq) is widely used to investigate chromatin accessibility but experiments often contain empty droplets, droplets containing damaged nuclei or cellular-debris. Accurately filtering high-quality nuclei is important as noisy droplets contribute reads that can distort downstream analysis and obscure rare cell population. Current approaches often rely on read depth and or predefined genomic features that are not exactly the same as chromatin accessibility landscape. Here, we present a Shannon entropy-based method that filters high-quality nuclei more robustly by quantifying genome-wide accessibility complexity. This approach improves signal-to-noise ratio, increases sensitivity for accessible-region detection, and enables more reliable downstream inference in scATAC-seq studies.

## Background

In eukaryotes, DNA is densely compacted into chromosomes by wrapping around core histones to form nucleosomes [1], which are then hierarchically folded into higher-order chromatin structures. Nucleosome density controls the accessibility of genomic loci to be machinery responsible for functional regulatory processes. Open chromatin regions, where nucleosomes are loosely packed, allow the corresponding DNA sites to interact with regulatory proteins such as transcription factor which can initiate transcription. Whereas closed chromatin regions, where nucleosomes are tightly packed, usually corresponds with gene silencing or suppression. Analyzing chromatin accessibility in different cell-types and cell-states helps us better understand gene regulation. Single-cell Assay for Transposase-Accessible Chromatin by sequencing (scATAC-seq) has become a widely used method for investigating chromatin accessibility with single-cell resolution. This method uses transposases enzyme Tn5 to insert sequencing adapters into DNA at accessible chromatin regions within each nucleus for tagmentation [2]. DNAs are then fragmented and barcoded at tagmentation sites for amplification and sequencing. Sequencing reads are aligned back to the reference genome and the accessibility of genomic intervals is inferred from read count for each cell, represented by a unique barcode sequence.

Because scATAC-seq data is sparse [3], single-cell libraries are aggregated to generate an ensemble library that is used for inference of accessibility peaks across the entire cellular population. Subsequently, individual cells are assigned feature values from the ensemble peak set to identify distinct epigenetic states. However, various sources of experimental artifacts can yield low quality single-cell observations that hinder peak inference and pollute the population measurements. These artifacts include background reads that come from empty droplets [4], or fragments from partially digested nuclei or dead cells [5] that lost their intact 3D genome structure. Reads associated with noisy droplet barcodes containing these cellular artifacts can impair downstream inference by elevating background noise and obscuring true accessibility signals. Because peak-calling algorithms identify peaks from local enrichment of sequencing reads [6–11], spurious reads reduce both the sensitivity and accuracy of peak detection. Additionally, these low-quality barcodes can interfere with cluster identification. Therefore, accurately identifying high-quality nuclei barcodes and filtering out barcodes associated with empty droplets or low-quality nuclei is a key pre-processing step for reliable downstream analyses.

There are two commonly used approaches to filter high-quality barcodes in scATAC-seq experiments. One approach, implemented in CellRanger-arc [12], first constructs a pseudo-bulk peak set after aggregating all reads in the experiment, then subsequently evaluates each barcode for the enrichment of its reads within these peaks. Because the reference peak set is dominated by signals from abundant cell populations, this approach may fail to detect low abundance cell types in heterogeneous samples [4]. In addition, droplets containing ambient DNA fragments from highly accessible regions, such as housekeeping genes, may exhibit apparent enrichment within the pseudo-bulk peaks and be consequently misclassified as high-quality barcodes. Another approach, such as in [9–11], evaluates barcode quality by measuring enrichment of reads in predefined genomic feature regions, most commonly transcription starting sites (TSS), to distinguish barcodes associated with high-quality nuclei. Because these methods rely on TSS enrichment (TSSe) score as a surrogate measure of chromatin accessibility, their performance depends on the assumption that TSS regions are representative of overall quality of chromatin accessibility. Moreover, the TSSe score is sensitive to both experimental factors (e.g., the relative Tn5 concentration to genomic DNA [5]), and differences in computational implementations choices [13], including the size of the flanking region around the TSS, potentially affecting the robustness and consistency of quality assessment. In addition, while these approaches are powerful and widely used, neither of these approaches can be used alone to filter high-quality nuclei - they have to be accompanied by other criteria such as total number of fragments, which is also highly dependent on experimental design.

To address these limitations, we developed a principled framework to quality control in scATAC-seq, termed SADE (Shannon entropy for scATAC-seq denoising). SADE quantifies the Shannon entropy of the distribution of genomic insertion sites within each barcode’s fragment library, thereby using the complexity of chromatin accessibility patterns as an indicator of nucleus quality. This enables cell-type-agnostic identification of high-quality nuclei while filtering out barcodes associated with droplets containing ambient DNA, degraded chromatin, or cellular-debris with mixed chromatin fragments.

Here, we demonstrate that SADE more effectively identifies noisy droplets containing degraded DNA or anomalously high fragment counts. We further show that performing quality control with SADE improves downstream analysis by increasing the sensitivity of peak inference in scATAC-seq and enhancing the recovery of high-quality RNA profiles in paired multiome (ATAC+GEX) data.

## Results

### SADE: Method Overview and Motivation

Chromatin accessibility is highly heterogeneous across the genome because the nucleosome occupancy and positioning are dynamically regulated across promoters, enhancers, insulators and other functional genomic regions [14]. Consequently, Tn5 insertions in scATAC-seq data are expected to exhibit complex yet highly structured genome-wide accessibility patterns that reflect the underlying chromatin architecture. In contrast, empty droplets, and droplets containing degraded nuclei or cellular-debris lack this organized accessibility landscape and therefore produce sparse, noisy, or otherwise aberrant insertion patterns (**Fig 1A**). This biological distinction motivates the use of Shannon entropy to quantify the complexity of genome-wide Tn5 insertion patterns as a measure of nucleus quality. Based on this principle, we developed SADE, a quality control framework for scATAC-seq that models the Shannon entropy of each barcode’s genome-wide insertion profile to identify high-quality nuclei. Intact nuclei are expected to exhibit characteristic entropy profiles arising from their structured chromatin accessibility landscape, whereas technical artifacts yield abnormal entropy values. By automatically identifying distributional inflection points in the entropy distribution across barcodes, SADE provides a fully data-driven approach for distinguishing biological informative nuclei from technical artifacts. An overview of the SADE workflow is illustrated (**Fig 1**).

**Fig 1.**
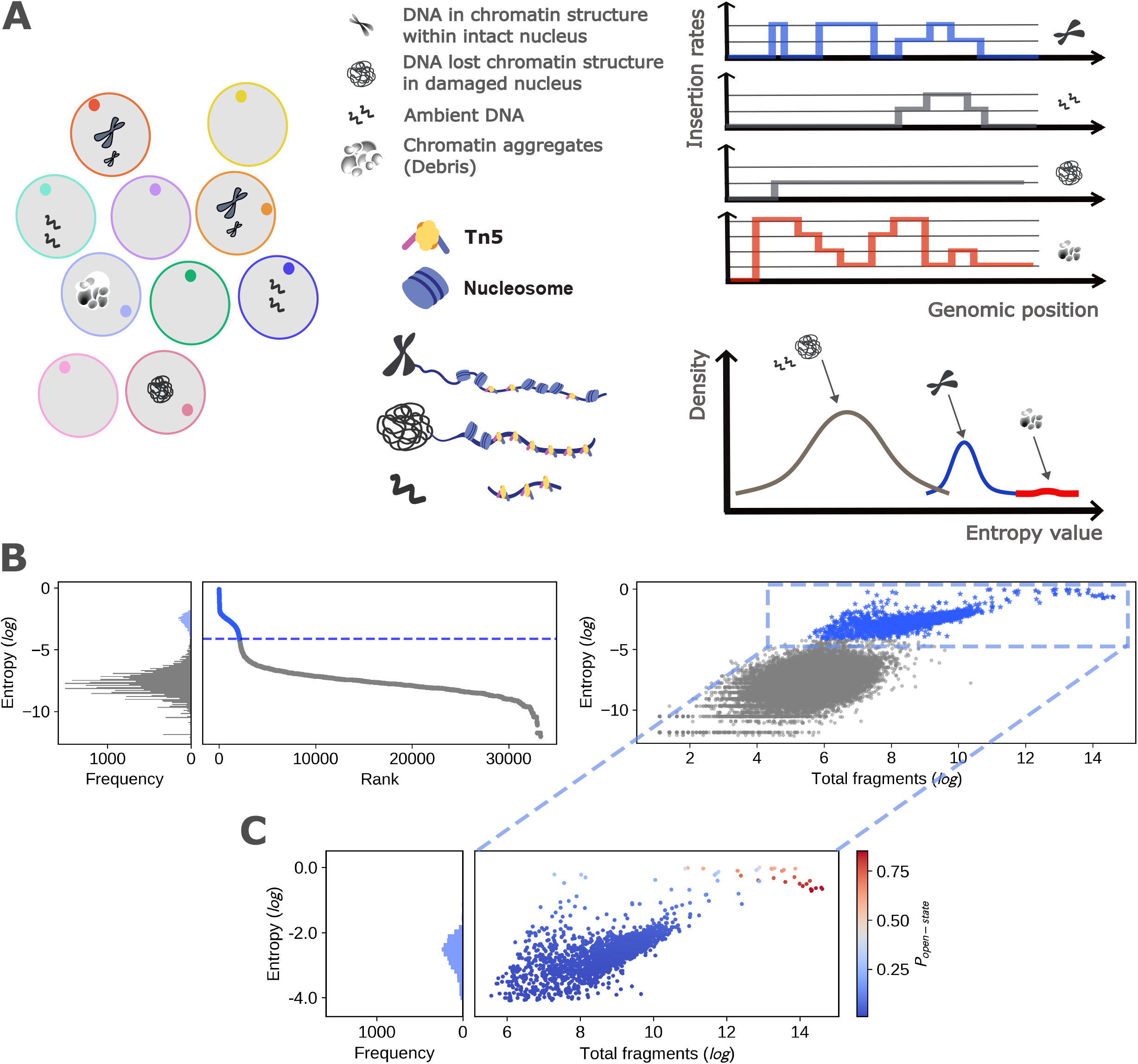
Method overview. A. **Left panel**: schematic of droplet types in a scATAC-seq experiment with individual droplet associated with a unique barcode sequence bead: (1) droplets containing a nucleus in its native chromatin state, (2) droplets containing a damaged nucleus with partially disrupted chromatin structure, (3) droplets containing ambient DNA fragments, and (4) droplets containing cellular debris represented by chromatin aggregates. **Right panel**: Schematic of Tn5 insertion patterns across the genome for each droplet type (**top**) and the corresponding genome-wide insertion complexity, measured by entropy (**bottom**). B. Schematic of SADE’s first step using mixture distribution on entropy values to identify barcodes associated with empty droplets and droplets containing denatured nuclei with damaged chromatin. C. Schematic of SADE’s second step using estimated accessible genome fraction to identify barcodes associated with cellular-debris-containing droplets among those that passed automatic entropy value thresholding.

SADE partitions the reference genome into fixed-size windows and estimates the Tn5 insertion rate within each window using the ends of sequencing reads. Windows with high insertion rates correspond to highly accessible chromatin sites, whereas windows with low insertion rates correspond to less accessible sites. SADE then quantifies the complexity and heterogeneity of the genome-wide insertion-rate distribution for each barcode using Shannon entropy. Entropy value is then normalized by read depth within each barcode to facilitate robust detection of high-quality nuclei across barcodes with sufficient read depth (**Methods**). The entropy values of all barcodes in an experiment form a mixture population where barcodes associated with intact nuclei concentrate at high entropy values and low-quality droplets concentrate at low entropy values. To distinguish them, a cubic spline is fitted to a curve of the ranked entropy values, and its first and second derivatives are calculated to identify the density inflection point. This inflection point serves as the optimal cutoff; barcodes with entropy values below this threshold are detected as empty droplets containing ambient DNA fragments or those with damaged chromatin (**Fig 1B**). SADE takes the post-alignment fragment file and calculates Shannon entropy value for each barcode under a mixture distribution. The mixture distribution contains a Poisson distribution to model accessible regions’ insertion rates, and a Dirac delta function at zero to model closed regions, with mixture rate inferred from data. The inferred mixture rate represents genome fraction that is accessible 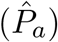 or closed 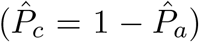. Low-quality droplets are filtered in two steps. First, barcodes associated with empty droplets or damaged nucleus corresponding to low entropy are filtered out with automatic inflection point thresholding on entropy values. We then further filter out barcodes associated with droplets containing cellular-debris which have accessible genome fraction that exceeds the ATAC assay’s signal coverage’s full potential by setting a threshold at 0.5 as default (**Fig 1B, Method**). Those droplet barcodes tend to have greater read-depth and outlier high entropy values (**Fig 1C**, **Methods**).

### SADE detects chromosome abnormalities overlooked by other filtering criteria

To confirm that barcode filtering using SADE produces reasonable single-nuclei libraries, we analyzed ATAC fragments from a public human multiome (ATAC+GEX) PBMC benchmarking dataset [15]. SADE identified 2,101 high-quality nuclei barcodes across a read-depth range of 261 to 398,925, showing high concordance with other multi-metric filtering approaches (**Fig 2A, Methods**). Specifically, 1,865 (88.77%) of the SADE-identified barcodes were also classified as high-quality by CellRanger’s criteria—which uses the total number of reads overlapping with ensemble peaks—and 1,910 (90.91%) barcodes overlapped with ArchR’s criteria, which utilizes TSSe scores alongside read depth (**Sup fig1A**).

**Fig 2.**
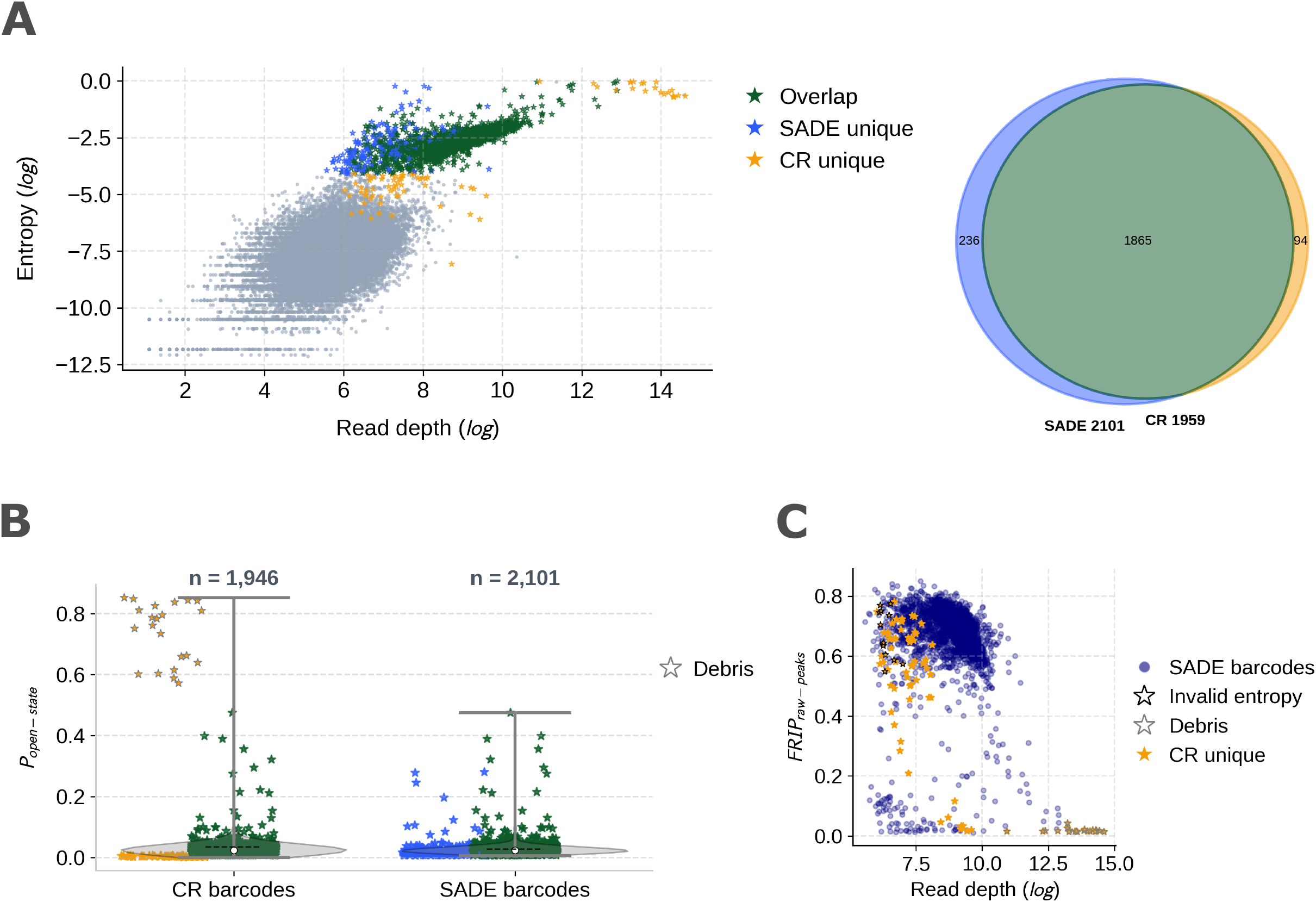
Shannon entropy measured complexity of genomic accessibility allows for detection of noisy droplets. A. **Left panel**: scatter plot of entropy versus ATAC fragment read depth in the PBMC dataset. Each dot represents a droplet barcode; star-shaped dots indicate the 2195 barcodes classified as high-quality nuclei by SADE and/or CellRanger, colored green for agreement between both methods, blue for SADE only, and yellow for CellRanger only. Circular dots represent barcodes classified as noisy droplets by both methods. **Right panel**: Venn diagram showing agreement and disagreement among the 2195 filtered barcodes identified by SADE and CellRanger. Numbers in each region indicate the number of barcodes in each intersection. Both panels use the same color scheme. B. Violin plot showing the distribution of SADE-estimated open-state probability *P*_*a*_ (i.e., accessible genome fraction) across barcode sets permitted by each method. Each star-shaped dot represents a barcode and is colored by the method that classified it as high quality. Star-shaped dots with gray outlines were flagged by SADE as DNA debris with *P*_*a*_ *>* 0.5. C. Scatter plot of the fraction of reads in peaks versus ATAC fragment read depth for the 2195 barcodes retained by SADE and/or CellRanger. Blue circular dots denote SADE-identified high-quality nuclei. Yellow star-shaped dots indicate the 94 barcodes uniquely admitted by CellRanger. Star-shaped dots with black outlines were barcodes classified by SADE as low-complexity chromatin accessibility with invalid entropy values.

Within the union set of 2,384 barcodes classified as high quality by at least one of the three methods, SADE uniquely identified 118 barcodes that had lower overall read depths. This allows for the possibility of recovering high-quality nuclei with low read depths, whereas the other two approaches utilize read depth as a filtering criterion, either implicitly or explicitly. Additionally, SADE flagged 22 CellRanger-permitted barcodes with excessively high read depths as cellular-debris, by thresholding estimated accessible genome fraction at 0.5 (**Fig 2B, Sup fig1B, Methods**). At the individual barcode level, these barcodes associated with cellular-debris-containing droplets consistently exhibited higher read depth across the entire genome than a diploid cell, which has a maximum of 2 copies, suggesting they represent chromatin aggregates from multicellular content (**Sup fig2A, Sup fig2B**). We calculated the fraction of reads overlapping with peaks (FRIP) by first inferring raw peaks from all reads and found that cellular-debris-containing droplets had extremely low FRIP scores (**Fig 2C, Methods**). While CellRanger and SADE nuclei sets exhibited comparable FRIP score distributions, with averages of 65.41% and 64.82% respectively (**Sup fig1C, Methods**), barcodes filtered uniquely by ArchR TSSe all showed low FRIP scores with highest at only 10.62%. Among the CellRanger-permitted barcodes, SADE flagged 13 barcodes with invalid entropy due to average Tn5 insertion rates being too low to infer genomic accessibility complexity, despite their high FRIP scores (**Fig 2C, Methods**). An additional 8 barcodes admitted uniquely by ArchR TSSe were also marked as invalid. Single-barcode read-coverage analysis showed a maximum read depth of only 1 across entire chromosomes in these invalid-entropy droplets, suggesting chromosomal loss in these human PBMC cells (**Sup fig3A, Sup fig3B**). Similar to fragment length distribution (FLD) patterns from cellular-debris droplets, some barcodes associated with invalid-entropy droplets displayed normal multi-peak patterns (**Sup fig 2C, Sup fig3C**), indicating that damaged nuclei with chromosomal loss can be easily overlooked when relying on FRIP score or FLD filtering.

### Removing low-quality barcodes improves downstream peak detection sensitivity

We hypothesized that a principled and unbiased read filtering approach could improve peak calling by systematically removing droplets with contaminating reads. For example the 22 barcodes identified as cellular-debris-containing droplets by SADE but retained by CellRanger in the analysis above contributed 22.14 million DNA fragments alone, of the 43.29 million reads across 1959 barcodes, while SADE identified 2101 high-quality nuclei barcodes containing a substantially lower total of 21.45 million reads (**Fig 3A, Sup fig4A**). Importantly, the reads from cellular-debris-containing droplets were nearly uniformly distributed across the genome, despite having fragment lengths enriched at the mononucleosome size in FLD (**Fig 3B, Fig 3A, Sup fig2C**). This observation implies that filtering these reads before peak calling might improve peak detection.

**Fig 3.**
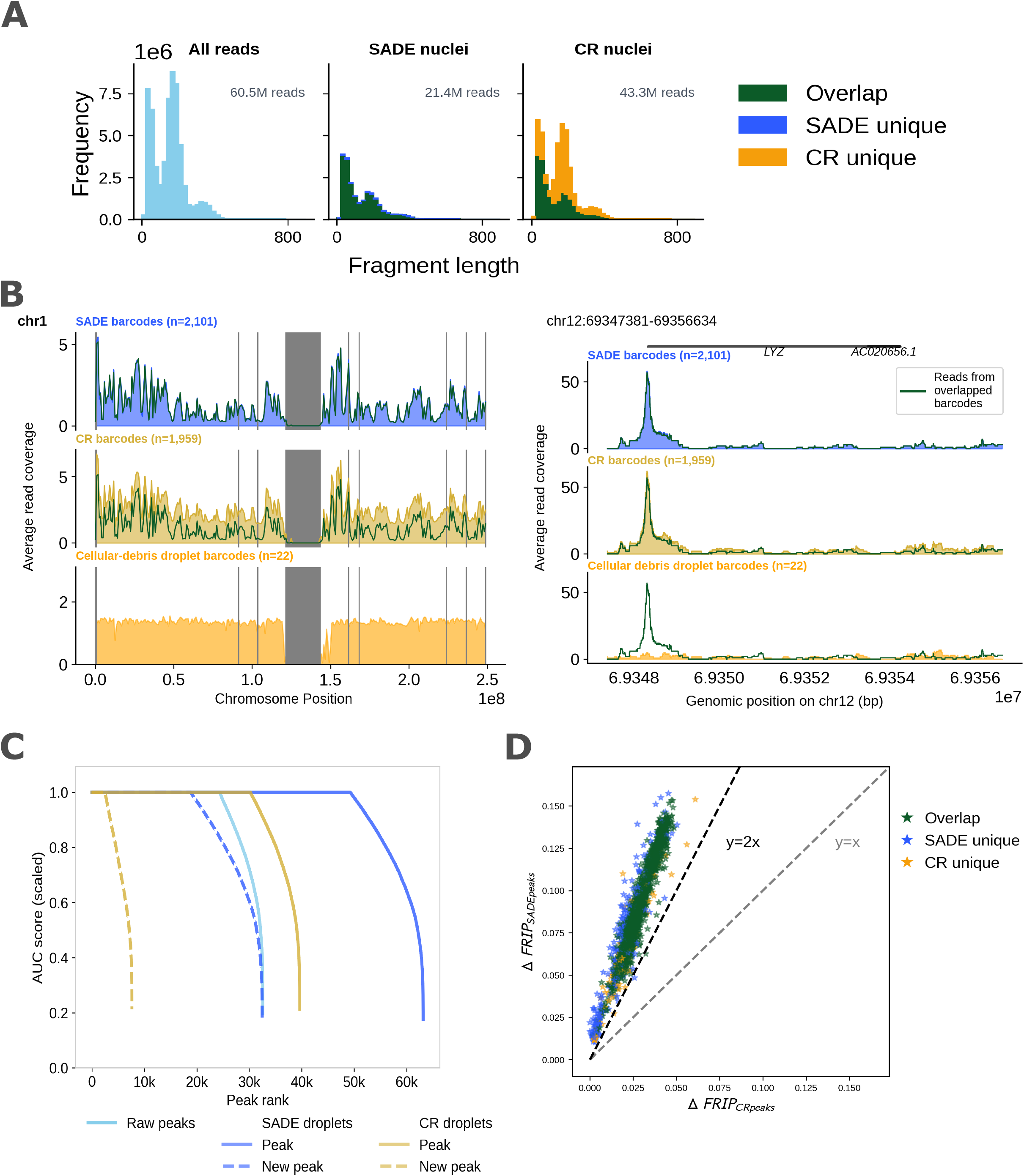
High quality barcode filtering improves downstream detection sensitivity of accessible regions. A. Fragment length distribution of reads from each barcode set: the first column shows all reads in the experiment; the other two show reads from barcodes retained by each method. Reads were colored at droplet barcode level with colors corresponding to methods that determined them as high quality. B. Normalized read coverage at selected regions. The top row shows barcodes retained by SADE, the second row those retained by CellRanger, and the third row the CellRanger-retained subset flagged as cellular-debris by SADE. Reads in shared barcodes are shown in green line for reference in the top two rows on the left panel and in all rows on the right panel. Blacklist regions are shaded grey. **Left panel**: across whole chr1. **Right panel**: LYZ gene region. C. Ranked peaks by area under the curve (AUC) score for each peak set. Higher AUC indicates stronger read pile-up in the pseudo-bulk profile. Raw peaks (sky blue) were inferred from all reads, while method-specific peak sets (SADE, blue; CellRanger, yellow) were inferred from method-retained barcodes only. Newly discovered peaks were identified by intersecting each method-specific set with raw peaks. D. FRIP score increase for each droplet in the 2195 barcodes, colored by the method retaining it. The x-axis shows the FRIP difference between using CellRanger peaks and raw peaks, the y-axis shows the FRIP difference between using SADE peaks and raw peaks. Reference lines Y=X and Y=2X are shown.

To assess how low-quality nuclei impacts downstream peak feature inference, we used Genrich [6] to infer peaks using only the reads from filtered barcodes, permitted by each filtering method and compared the resulting peak sets to the raw peaks inferred from unfiltered ensemble reads. Compared to the 32,572 raw peaks detected using all reads, SADE-filtered reads enabled the detection of 32,461 new sites (99.66% increase) that did not overlap with the raw peaks (**Fig 3C**). In contrast, CellRanger nuclei only detected 7,622 new sites (23.40% increase), reflecting its higher background noise and reduced signal-to-noise ratio (**Fig 3C, Methods**). The new peaks had shorter genomic length, demonstrating that removing noisy reads significantly increases genomic accessibility signal resolution in a cell-type agnostic manner (**Sup fig4B**).

Recalculating the FRIP score with these higher-resolution peak sets, we observed a universal increase in all barcodes compared to that using raw peaks. However, the increase in FRIP score using the SADE peak set was, on average, more than two-fold greater than the increase observed using the CellRanger peak set (**Fig 3D**).

Similarly to SADE, the ArchR TSSe filtering approach allowed for the detection of 36,472 new sites (111.97%) using 18.84 million reads across 2,132 barcodes, resulting in a similar FRIP score increase across all union barcodes (**Sup fig4A, Sup fig4B**). Nonetheless, barcodes uniquely identified by ArchR TSSe consistently had the lowest fraction of reads overlapping any peak sets (**Sup fig4C, Sup fig4D**). This suggests that reads from barcodes that were uniquely permitted by ArchR TSSe may not contain the whole genome DNA but rather large genomic segments that happen to span highly accessible regions.

To see if these peaks overlap with highly accessible genomic regions, we annotate them with candidate cis-regulatory elements (cCREs) using the 2.3 million registered human cCRES in ENCODE [16], and found that all three filtering approaches enable peak detections in the high chromatin accessibility regions. Across approaches, over 99% of newly discovered peaks overlapped cCREs with high chromatin accessibility signals (**Sup fig4E, Methods**). We also observed that wider peaks tend to overlap more cCREs (**Sup fig4F, Methods**).

### Barcode filtering with SADE preserves cell clusters identified with scRNA-seq

To evaluate each scATAC-seq filtering method’s accuracy at the cell-type level, we independently processed, clustered, and annotated cell types using canonical marker genes [17, 18] from the RNA modality of the multi-ome PBMC measurements (**Fig 4A, Sup fig5, Methods**). The majority (1,760) of barcodes showing complete concordance across all three filtering methods were distributed among all existing RNA-annotated cell types. Conversely, within the 589 barcodes with any filtering disagreement, 314 of them exhibited low RNA signals and clustered into four noisy RNA clusters (c4, c9, c10, and c11) with similar gene expression profiles (**Fig 4A, 4B, Sup fig6A**). Specifically, scRNA-seq clusters c4, c9, and c10 were enriched with barcodes that failed RNA quality control, while c11 was enriched with ATAC cellular-debris-containing droplets (**Fig 4C, Sup fig6B, Methods**). Recognizing the systematic differences between RNA and ATAC modalities—arising from both biological factors (e.g., quiescent cells) and technological assay differences [19–22]—we thoroughly analyzed the scATAC-seq profiles to bridge this gap.

**Fig 4.**
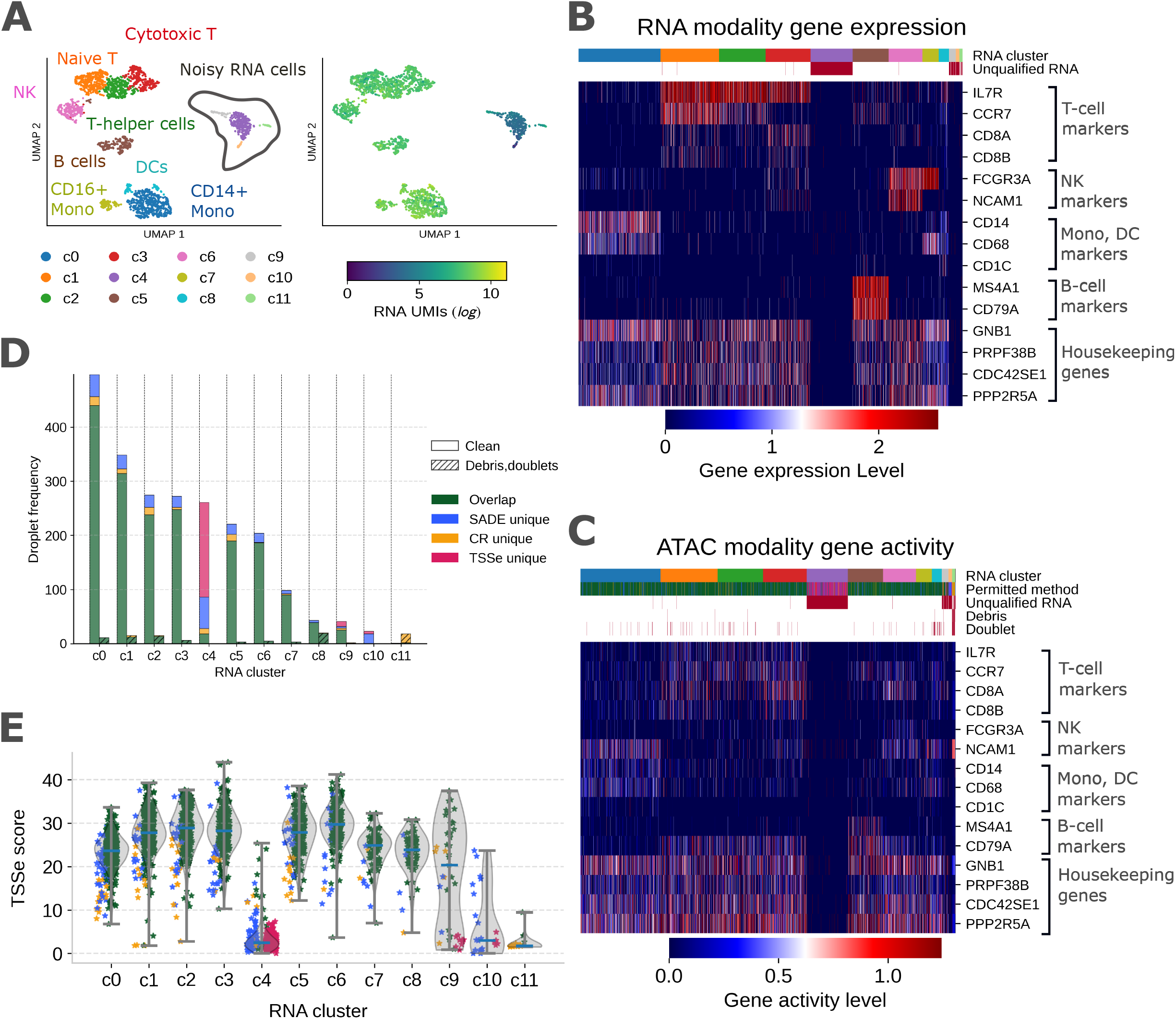
SADE identified high quality barcodes contain reads that have enriched signals in biological regions. A. Low-dimension representation UMAP of RNA modality gene expression for the 2381 union barcodes, annotated by cell type at the cluster level. Three of the 2384 barcodes lacked RNA profiles and are therefore excluded. Each dot represents a barcode and is colored by RNA cluster (**left panel**) or RNA UMI count (**right panel**). Circled noisy-RNA clusters predominantly comprise barcodes whose RNA profiles failed CellRanger’s RNA-filtering criteria. B. Heatmap of gene expression for the 2381 union barcodes, showing selected marker and housekeeping genes. Each column represents a barcode, colored by RNA cluster and highlighted in red for barcodes with unqualified RNA profiles. C. Heatmap of gene activity scores for selected marker and house-keeping genes across the 2384 union barcodes. Gene activity was computed using ATAC profiles. Each column represents a barcode and is colored by RNA cluster and ATAC filtering methods that retained it; barcodes with unqualified RNA profiles or unclean ATAC profiles (cellular-debris or doublets) are highlighted in red. D. Stacked bar plot showing the number of barcodes in each RNA cluster, colored by the ATAC filtering methods that retained each barcode. Within each cluster, barcodes are grouped by ATAC profile type (clean, doublet or cellular-debris). E. Violin plot showing distribution of TSSe scores calculated by Sna-pATAC within each RNA cluster and color coded by the ATAC filtering methods that retained the barcode. The same color scheme is used across all plots for ATAC filtering methods that retained the barcodes.

Analysis of the scATAC-seq profiles revealed 96 estimated doublets (4.3%) using AMULET [23]. These doublets were predominantly found in the common barcodes across all identified cell types (65 barcodes) or in the cellulardebris-enriched noisy scRNA-seq cluster c11 (14 barcodes) (**Fig 4C, 4D, Methods**). The noisy scRNA-seq clusters c4 and c10 (primarily consisting of 184 ArchR TSSe uniquely or 66 SADE uniquely permitted barcodes) and c11 (primarily from CellRanger permitted cellular-debris-containing droplet barcodes) all exhibited low TSSe scores (**Fig 4E, Methods**). Notably, the average TSSe score calculated by SnapATAC for ArchR TSSe uniquely identified barcodes was 2.62 (**Fig 4E, Sup fig6C**), which falls below the ArchR TSSe cutoff of 4, reinforcing the reported inconsistency in TSSe score calculations across different computational implementations [13]. Overall, the gene activity calculated using the ATAC modality closely resembles the gene expression observed in the RNA modality at the RNA cluster level (**Fig 4C, Methods**).

To further investigate epigenetic states, we clustered the scATAC-seq profiles using the 2,286 clean barcodes after removing cellular-debris-containing droplets and doublets from the union barcode set (**Fig 5A**). The clustering results showed high consistency between the ATAC and RNA modalities at the cell-type level (**Fig 5B, Sup fig7A**). Specifically, scATAC-seq cluster c0 (292 barcodes) was predominantly composed of nuclei from the noisy scRNA-seq clusters c4 (95.79%), c10 (73.91%), and c9 (34.88%), exhibiting low TSSe scores in ATAC modality and low read counts in both modalities (**Fig 5B, 5C, Sup fig7B**). Importantly, the lack of activity in DNA regions associated with housekeeping genes and quiescence markers suggests that these barcodes are likely noisy droplets incorrectly recognized as quality-nuclei by the filtering methods, rather than a rare cell population (**Fig 5D**). For SADE, permitting 91 such barcodes is due to tail overlap at the entropy distribution between the high-entropy and low-entropy populations; consequently, a small number of noisy droplets may still possess entropy values slightly exceeding the density inflection point used to separate the two populations. However, those falsely admitted barcodes exhibit cluster-level low entropy values (**Sup fig7C**) and thus can be removed during downstream cluster-level analysis.

**Fig 5.**
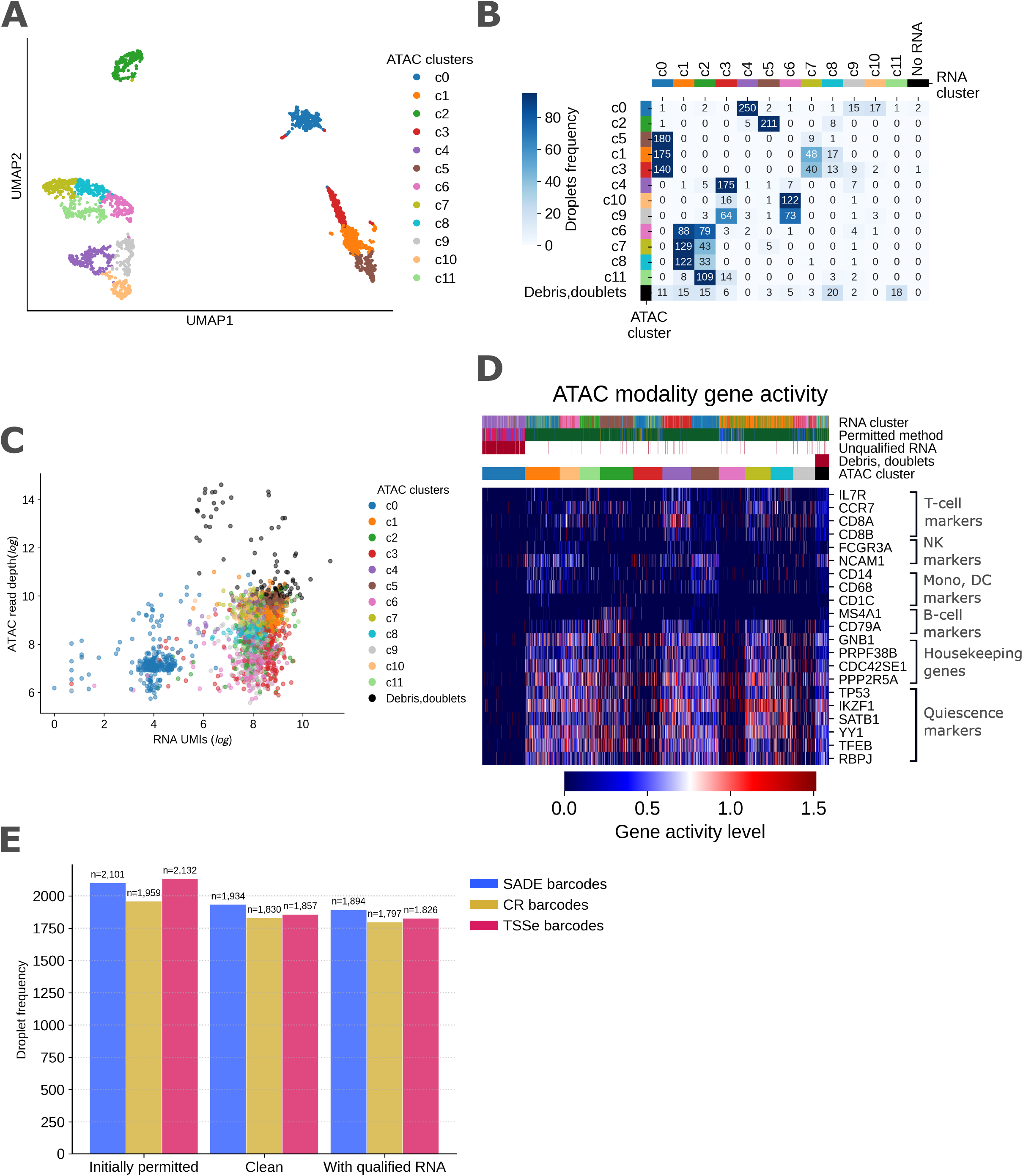
SADE assists fine resolution cell clustering and doublet detection. A. Low-dimension representation UMAP of ATAC profiles of the 2286 barcodes associated with clean nuclei, colored by ATAC clusters. B. Heatmap of droplet barcode frequency in each ATAC cluster– RNA cluster pair. C. Scatter plot of ATAC read depth against RNA UMI count for the 2381 union barcodes, color coded by ATAC cluster or in black as cellular-debris or doublets. D. Heatmap of gene activity scores for the 2384 union barcodes across selected marker, housekeeping, and quiescent-marker genes. Gene activity was computed from ATAC profiles. Each column represents a barcode and is colored by ATAC cluster, RNA cluster, and the filtering methods that retained it; barcodes with unqualified RNA profiles or unclean ATAC profiles (cellular debris or doublets) are highlighted in red. E. Bar plot showing barcode frequency in each filtering method at (1) initial stage, (2) after removing cellular-debris or doublets, and (3) last stage with qualified RNA profiles.

Taken together the above observations suggest that SADE outperformed both ArchR TSSe and CellRanger in the ability to filter high-quality nuclei. SADE identified the highest number of clean quality nuclei (1,934 barcodes) free from cellular-debris, doublets, or inefficient signals, surpassing ArchR TSSe (1,857 barcodes) and CellRanger (1,830 barcodes) (**Fig 5E**). Furthermore, SADE achieved the highest recovery of quality RNA profiles among these high-quality, clean scATAC-seq nuclei barcodes (**Fig 5E**).

## Discussion

We have developed a novel, cell type-agnostic method for barcode filtering in scATAC-seq experiments called SADE, which utilizes Shannon entropy under mixture-distribution modeling to quantify the complexity of genomic accessibility signals at the droplet barcode level. This principled approach reduces scATAC-seq experimental noise, by removing empty droplets with spurious fragments of DNA, and droplets containing damaged nuclei with digested chromatin or cellular-debris with abnormally high read counts.

In our PBMC multiome (ATAC+GEX) dataset benchmarking, SADE identified more clean nuclei than read-depth-based filtering strategies and outperformed TSSe- and CellRanger-based approaches in removing cellular-debris-containing droplets and doublets. It also recovered the highest number of barcodes with corresponding RNA profiles, supporting its ability to preserve biologically meaningful signal while excluding noise. These improvements translated into more reliable cell-type clustering and stronger agreement with matched RNA measurements. Across four additional datasets from human and mouse tissues, including a human 10k PBMC dataset, a human brain 3k dataset, a mouse brain 5k dataset and a mouse cryo-preserved brain 5k dataset, SADE showed consistent performance (**Sup fig8, Methods**), indicating broad applicability across species and tissue types. A limitation of the mixture-distribution approach is that a small fraction of low-quality barcodes may overlap with the high-entropy tail of the high-quality population, although these can often be removed in later cluster-level analyses.

We also evaluated parameter robustness and found that the default 3-kb genome bin size – a value determined by the average nucleosome cluster size [14] – provides stable separation between low- and high-quality barcodes across a wide range of bin sizes (**Sup fig9, Methods**). Performance remained consistent for window sizes between 500 bp and 10,000 bp in both human and mouse datasets, supporting the practical robustness of the method (**Sup fig9**).

In summary, SADE provides an unbiased and robust approach for barcode filtering in scATAC-seq using Shannon entropy, improving the identification of high-quality nuclei and reducing contamination from empty droplets, damaged nuclei, and cellular-debris. By enabling more accurate downstream analysis and better concordance with matched RNA data, SADE improves the reliability and interpretability of single-cell chromatin accessibility studies. The method is designed for broad use across datasets, species, and tissue types. SADE is available at https://github.com/streetslab_SADE.

## Methods

### Entropy Calculation for Individual Droplet Barcodes

For each droplet barcode, SADE models the genome-wide distribution of Tn5 insertions using a two-component mixture model. The reference genome is partitioned into *N* non-overlapping windows of fixed size (default 3-kb), where 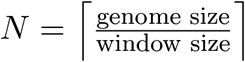 .

For the *i*-th genomic window (*i* = 1, …, *N* ), let *Y*_*i*_ denote its latent accessibility state, where *Y*_*i*_ = 1 indicates that the window is accessible and *Y*_*i*_ = 0 indicates that it is inaccessible. Let *X*_*i*_ denote the number of Tn5 insertions within that window. For barcode *b ∈ {*1, …, *B}*, where *B* being the total number of barcodes, we assume the hierarchical model

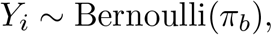

and

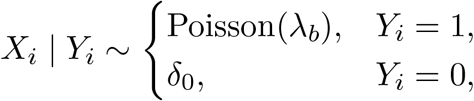

where *π*_*b*_ denotes the proportion of accessible genomic windows for barcode *b, λ*_*b*_ (*λ*_*b*_ *>* 0) is the expected Tn5 insertion rate within accessible windows, and *δ*_0_ denotes a point mass at zero corresponding to inaccessible windows. The barcode-specific parameter *π*_*b*_ reflects the overall accessibility of the chromatin and is therefore informative of nucleus quality, whereas *λ*_*b*_ accounts for barcode-specific sequencing depth variation.

After fitting the above mixture model, SADE uses the estimated genome-wide insertion-rate distribution for each barcode to compute its Shannon entropy, which serves as a quantitative measure of the complexity of the underlying chromatin accessibility landscape. The details of parameter estimation and entropy calculation are described below.

#### Estimation of *λ*_*b*_

Because only genomic windows with nonzero Tn5 insertion counts are observed from the Poisson component of the mixture model, we estimate the Poisson rate parameter *λ* using a zero-truncated Poisson distribution (ZTPD) and subsequently impute the zero-insertion windows.

Specifically, for the droplet with barcode *b*, let *M*_*b*_ denote the number of genomic windows (out of the total *N* windows) with nonzero Tn5 insertions, and let *{b*_*j*_; *j* = 1, …, *M*_*b*_*}* denote the indices of these windows. For each window *b*_*j*_, let 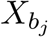 denote the observed number of Tn5 insertions and 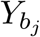 denote its latent accessibility state. Since 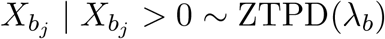 by our earlier assumption, the maximum likelihood estimator (MLE) 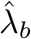 is obtained by maximizing the log-likelihood

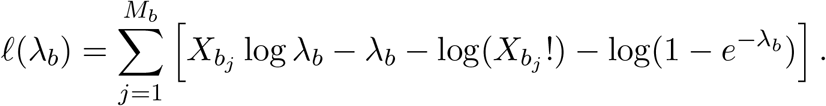

Let *R*_*b*_ denote the total number of Tn5 insertions for barcode *b*. Solving the likelihood equation using the principal branch of the Lambert *W* function [24], denoted by *W*_0_(*·*), yields the maximum likelihood estimator

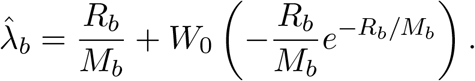

Since *R*_*b*_, *M*_*b*_ *∈* ℤ ^+^, the argument of the Lambert *W* function is always nonpositive. Moreover, the principal branch *W*_0_(*·*) is real-valued only for arguments greater than or equal to 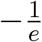, which is equivalent to 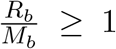. Note that the condition 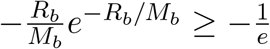 is guaranteed here since we have 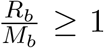. However, a strictly positive interior MLE for *λ*_*b*_ (*λ*_*b*_ *>* 0) requires 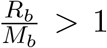. Therefore, SADE requires 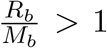. Barcodes that do not satisfy this condition are flagged as having invalid entropy estimates.

#### Estimating the Number of Zero-Insertion Accessible Windows

We can now estimate the number of accessible genomic windows with zero observed Tn5 insertions arising from stochastic sampling or insufficient sequencing depth. Let *M*_0_ denote the number of such unobserved windows. Under the fitted zero-truncated Poisson model, its estimate is given by

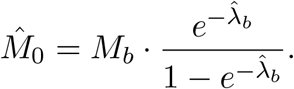

#### Estimating Accessible and Inaccessible Window Proportions

The estimated number of inaccessible genomic windows for barcode *b*, denoted by *M*_*b,c*_, is obtained by subtracting the estimated number of accessible windows from the total number of genomic windows. Specifically, 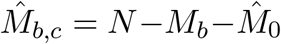, where *N* is the total number of genomic windows as defined before, *M*_*b*_ is the number of observed windows with nonzero Tn5 insertions, and 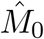 is the estimated number of accessible windows with zero observed insertions. Because *M*_*b,c*_ represents the number of inaccessible genomic windows, we require 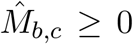. Barcodes violating this constraint are flagged as having invalid entropy estimates. Intuitively, a negative value of 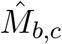 indicates that the fitted model imputes more zero-insertion accessible windows than can be accommodated by the observed data. This typically occurs when a large proportion of observed windows have very low Tn5 insertion counts, which may arise from over-tagmented nuclei or nuclei with disrupted chromatin architecture.

The estimated proportions of accessible and inaccessible genomic windows are then given by

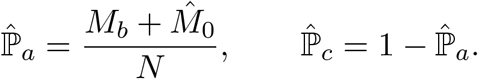

The two conditions leading to invalid entropy estimates can be combined into the single validity criterion 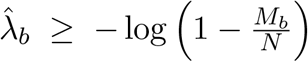 . Failure to satisfy this condition indicates that the estimated Tn5 insertion rate is too low to support the observed number of nonzero windows under the fitted mixture model. Such barcodes are flagged as having invalid entropy estimates.

#### Entropy Calculation

With the estimated proportions of accessible and inaccessible genomic windows, the Shannon entropy of the genome-wide Tn5 insertion process can be computed using the entropy decomposition formula

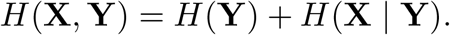

For barcode *b*, the estimated numbers of accessible and inaccessible genomic windows are *M*_*b*_ + 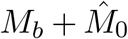 and 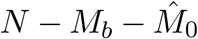, respectively. Therefore, assuming that *Y*_1_, …, *Y*_*N*_ are independent, we have

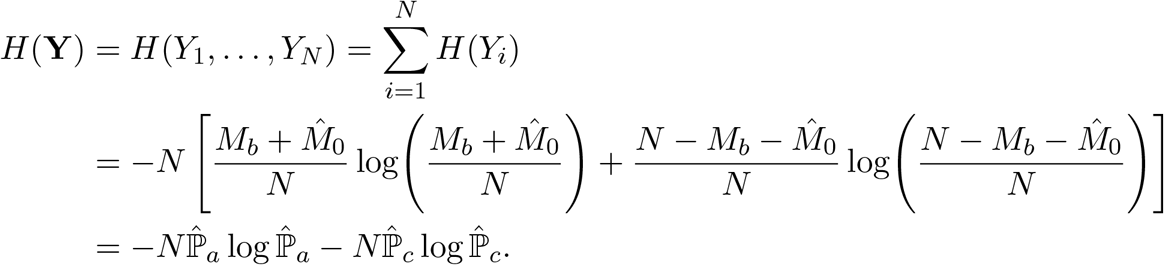

Under the conditional mixture model described earlier, the insertion counts *X*_1_, …, *X*_*N*_ are conditionally independent given *Y*_1_, …, *Y*_*N*_ . Consequently,

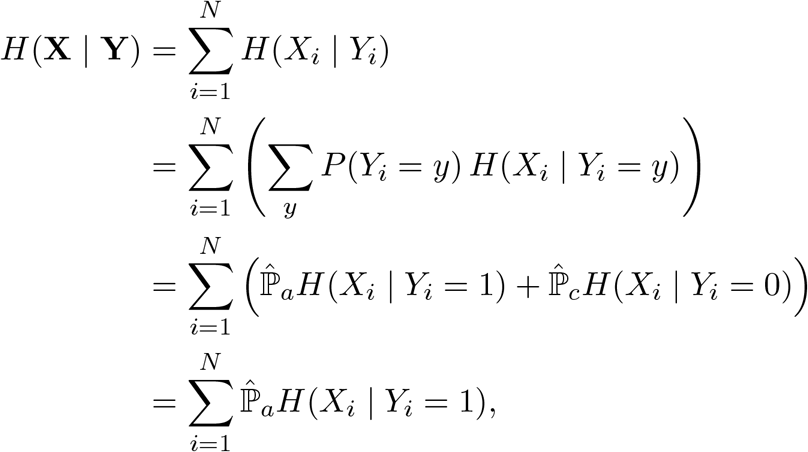

where the last equality follows from that *X*_*i*_ = 0 with probability one when *Y*_*i*_ = 0 and hence *H*(*X*_*i*_ | *Y*_*i*_ = 0) = 0. Furthermore, since *X*_*i*_ |*Y*_*i*_ = 1 ∼ Poisson*∼* 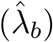, we have

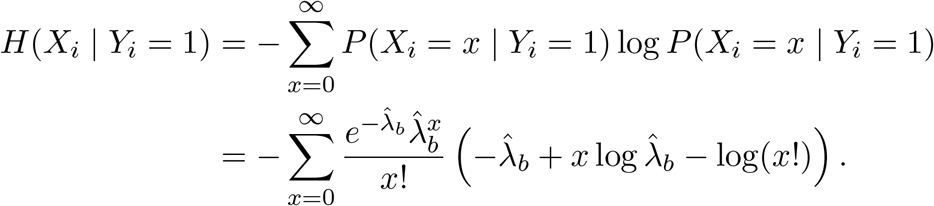

In practice, the infinite sum is truncated at the maximum observed insertion count, max *{x*_1_, …, *x*_*N*_*}*, for numerical computation, where (*x*_1_, …, *x*_*N*_ ) denote the observed Tn5 insertion counts across the *N* genomic windows for barcode *b*.

#### Sequencing-Depth Normalization

Because the entropy of the Tn5 insertion process depends on sequencing depth, we normalize the conditional entropy of accessible regions by the total number of observed Tn5 insertions to facilitate comparisons across droplets. Specifically,

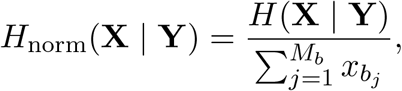

Where 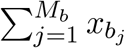 denotes the total number of observed Tn5 insertions for droplet with barcode *b*. The normalized entropy is computed independently for each droplet. Droplets that fail the validity criterion described above are assigned missing entropy values (i.e. invalid entropy values).

### SADE two-step filtering noisy droplet barcodes

In one ATAC-seq experiment, droplet-level entropy values calculated as stated above form a mixture distribution, where high entropy values correspond to barcodes associated with intact nuclei droplets, and low entropy values correspond to noisy droplets either with ambient DNA or a denatured nucleus with damaged chromatin structure.

A cubic spline is fitted to the curve of the ranked entropy values to separate these two populations. The first and second derivatives of the spline are calculated to identify local density inflection points, which serve as the candidates for the optimal population separation value.

*K* (default *k* = 2) inflection point candidates are chosen by highest entropy values. Among the candidates, the optimal inflection point is decided as the one with lowest 1st derivative values to minimize misclassification error.

Among those barcodes that passed the entropy threshold, they are further filtered using the estimated accessible genome fraction 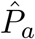 to remove barcodes that are putatively associated with cellular-debris. All the barcodes that have 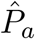 larger than predefined threshold (default at 0.5) are flagged as cellular-debris-containing droplets.

### Model robustness

#### Genome window size choice

The default genome window size is 3k-bp. Such a window size was chosen by qualitatively estimating the average nucleosome cluster size based on [14]. Specifically, nucleosome cluster sizes vary in genome DNA and is important for the spatial organization of chromatin structure. Cluster size of about 3-10 nucleosomes were shown in mammalian cells [25], as well as nucleosome free regions (NFRs). Cluster size of under 15 nucleosomes were observed in simulation under linker histon (LH) density of 0.5, short linker DNA lengths (LLs) 26-bp, and short NFRs 114-bp in the work of [14]. Given LH density of 0.5 was previously determined for mouse embryonic stem cells (mESC) [26], approximately 147-bp of DNA wrap around 8 core histone proteins to form a nucleosome, linker DNA lengths can span from 26-79 bp in Mesc [27] and NFRs are typically 80-400 bp, we want to have a window size that is on average able to cover a whole nucleosome cluster flanked by NFRs, we qualitatively set it to be 3k-bp.

In addition, a series of window size ( 100, 500, 1000, 3000, 6000, 10000, 15000, 20000, 30000, 50000, 80000, 100000, 200000, 400000, 800000-bp ) was examined. We found the separation of barcodes associated with quality nuclei droplets population from barcodes associated with noisy droplets population robust to window size between 500–10000-bp for genome sizes similar to human or mouse species (**Sup fig9**).

#### Chromosomes choice

The default chromosome used to calculate entropy is chromosome 1 (chr1). As entropy is calculated using the Tn5 insertions across all the windows under the mixture distribution SADE operates on, the number of windows can be regarded as sample size for a realization of the underlying Tn5 insertion distribution. We therefore chose chr1 – the largest chromosome in the genome, with the greatest number of insertion windows – to calculate entropy, because a larger sample size is expected to provide a better estimate of the underlying Tn5 insertion distribution.

Because it is unclear whether multiple chromosomes behave synchronously – for example, in cases involving chromosomal loss – we recommend using a single chromosome for SADE entropy calculation and barcode filtering. And we in general do not recommend using sex chromosomes for SADE barcode filtering.

SADE can, however, take multiple chromosomes for entropy calculation. In this case, the specified chromosomes are treated as one longer chromosome as if they have been virtually concatenated together (i.e., assuming they behave synchronously). This option is intended for species with very small genomes and extremely short individual chromosomes, where even the largest chromosome may contain too few Tn5 insertion windows to provide a sufficiently powered estimate.

For the reasons above, in our analysis, entropy values were calculated for the species’ largest chromosome chr1 rather than for the whole genome. We recommend for genome comparable in size to those of humans or mice, the largest chromosome should be used for SADE entropy calculation and filtering.

### Datasets

The main multi-ome PBMC dataset used in the manuscript is the MO V2 experiment from [15]. We downloaded its sequencing data from Sequence Read Archive under SRR24422953 [28] for ATAC fastq files, and SRR24446480 [29] for RNA fastq files.

The other datasets used in the manuscript are two mouse brain datasets [30, 31], two human dataset [32, 33] from 10x Genomics.

### Preprocessing scATACseq and scRNAseq sequencing reads

Sequencing data of the MO V2 multi-ome PBMC data were pre-processed independently for ATAC and RNA modality. Specifically, CellRanger-arc was used to pre-process ATAC sequencing fastq files [28] to get fragments.tsv.gz file for SADE to use as input to identify high quality nuclei barcodes as well as `possorted_bam.bam`for Genrich to infer peaks, and CellRanger was used to pre-process RNA sequence fastq files [29] to get gene expression count matrix `raw_feature_bc_matrix.h5` for analysis. Droplet barcodes were one-to-one mapped from ATAC and RNA whitelist droplet barcode lists at row index instructed by CellRanger-arc [34].

The reference genome used in sequence alignment for human cells is GRCh38, for mouse cells GRCm38.

### Comparison of barcode filtering among SADE, CellRangerarc, TSSe

SADE quality barcode filtering was obtained by running SADE on fragments.tsv.gz file. CellRanger-arc identified quality nuclei were directly obtained from filtered barcodes.tsv in the CellRanger-arc pre-processing result. TSSe nuclei filtering approach identified quality nuclei was obtained using ArchR with fragments.tsv.gz file and parameter setting at minimal TSSe score being 4 and minimal read depth as 1000. The three filtered droplet barcode sets were then used for comparison analysis.

### Peak detection, comparison, annotation

Genrich was used to infer peaks for each peak set with corresponding bam files. Specifically, raw peaks were inferred with `possorted_bam.bam` file, the peak set using reads in SADE nuclei was inferred with a subset of reads from this bam file by filtering on SADE admitted quality barcodes. And in the same way, the peak set using reads in CellRanger nuclei, the peak set using reads from ArchR-TSSe nuclei, as well as the peak set using reads from common nuclei were inferred. All peak inference with Genrich used the same parameter setting: removing PCR duplicates, with only reads that have quality measurement MAPQ >= 31, and at ATAC mode.

Each inferred peak set was then intersected using bedtools [35] with its species blacklist region [36] to remove peaks that fell in those regions. All other peak sets acquired using reads contained in its corresponding droplet barcodes were intersected with raw peaks using bedtools to annotate each peak as either newly-discovered, dropped, or recovered.

cCREs annotation to each peak was acquired by intersecting each peak with human cCREs class annotation (v4) downloaded from ENCODE SCREEN [37]using bedtools. In total, 8 cCREs classes were annotated in the downloaded human cCREs reference: Promoter-like signatures (PLS), proximal enhancer-like signatures (pELS), distal enhancer-like signatures (dELS), Chromatin accessibility + H3K4me3 (CA-H3K4me3), Chromatin accessibility + CTCF (CA-CTCF), Chromatin accessibility + transcription factor (CA-TF), Chromatin accessibility (CA), Transcription factor (TF). While all the other 7 classes correspond to high chromatin accessibility signals, TF class corresponds to low experimental chromatin accessibility cCREs regions.

### scRNAseq analysis

Multi-ome MO V2’s RNA gene expression count matrix was obtained in `raw_feature_bc_matrix.h5` from CellRanger RNA pre-processing result. Only RNA droplet barcodes that have correspondence ATAC droplet barcodes in the union 2384 barcodes were used for analysis. Among those 2384 ATAC union barcodes, 3 barcodes that were uniquely identified by SADE did not have corresponding RNA profiles, thus only 2381 barcodes with RNA profiles were used for analysis. These 2381 barcodes were further binary marked for their RNA profile quality by intersecting with RNA filtered droplet barcodes under `filtered_feature_bc_matrix`.

All 2381 droplets’ RNA profiles were used to cluster into 12 groups using SCVI [38]. Among them, 4 clusters (RNA c4, c9, c10, c11) have low gene expression signals. The other clusters were cell-typed with canonical markers (T cell markers: IL7R, CD3G, CD3E; naive T cells markers: CCR7; cytotoxic T cells markers: CD8A, CD8B; naive T helper cells markers: CD4; NK cells markers: FCGR3A, NCAM1; dendritic cells markers: CD1C, SERPINF1, HLA-DRA; monocytes markers: CD14, CD68, HLA-DRA).

Housekeeping genes used are: GNB1, PRPF38B, CDC42SE1, PPP2R5A; and quiescence markers used are: TP53, IKZF1, SATB1, YY1, TFEB, RBPJ.

### Analysis of scATACseq

The 2384 union ATAC droplet barcodes were used for downstream analysis. First doublet detection on scATACseq using AMULET [23]. Removing doublets estimated by AMULET as well as cellular-debris estimated by SADE in the union barcodes, the resulting 2286 barcodes were clustered using Snap-ATAC [11] with their corresponding fragments filtered from fragments.tsv.gz . SnapATAC clusters ATAC profiles with a tile matrix that was generated by binning the genome and counting fragments in each genome bin. 12 clusters were obtained at leiden clustering resolution at 1.5. Gene activity score was also calculated using SnapATAC for each of the barcode in the union set.

## Declarations

### Ethics approval and consent to participate

Not applicable.

### Consent for publication

Not applicable.

### Availability of data and materials

The five datasets analysed during the current study are all from publicly available data. The multi-ome PBMC dataset used in the manuscript is the MO V2 experiment from [15]. We downloaded its sequencing data from Sequence Read Archive under SRR24422953 [28] for ATAC fastq files, and SRR24446480 [29] for RNA fastq files. The other four datasets used in the manuscript are two mouse brain datasets [30, 31], two human dataset [32, 33] from 10x Genomics.

SADE software is available at https://github.com/streetslab/SADE. Code for the analysis is available at https://github.com/streetslab/SADE_manuscript.

### Authors’ contributions

SY and AS conceived the project. SY developed the method with input from AS, HH, KA, and SB. SY, JJL, and JG performed analysis. SY and KA wrote the software with input from JG. SY wrote the manuscript with input from all authors. HH provided guidance on the statistical modeling. AS supervised the study.

## Acknowledgements

We thank Rodrigo Cotrim Chaves for important discussions and suggestions during the early stage of this study. We also thank Mary Elizabeth Patti and Yu-Hua Tseng whose collaboration and discussion motivated the development of SADE.

