## Supplementary material for "Denoising single-cell ATAC-seq data with Shannon entropy": supplememtary figures and legends

### Supplementary figures and legends

October 21, 2026

**Sup fig 1.**

A. **Left panel**: scatter plot of entropy versus ATAC fragment read depth in the PBMC dataset. Each dot represents a barcode; star-shaped dots indicate the 2323 barcodes classified as high-quality nuclei by SADE and/or ArchR-TSSe, colored green for agreement between both methods, blue for SADE only, and pink for ArchR-TSSe only. Circular dots represent barcodes classified as noisy droplet by both methods. **Right panel**: venn diagram showing agreement and disagreement among filtered 2323 barcodes between SADE and ArchR-TSSe. Numbers in each section are the number of barcodes in each intersection.

B. Violin plot showing distribution of ATAC fragment read depth across filtering method identified quality-nuclei barcodes. Each star-shaped dot is a barcode and was colored by methods that determined them as high quality. Star-shaped dot with grey edge indicates SADE flagged cellular-debris.

C. Violin plot showing distribution of fraction of reads in raw peaks across each filtering method identified quality-nuclei barcodes, color coded by the method retained it.

D. **Left panel**: scatter plot of FRIP scores of all the 2384 barcodes that passed at least on filtering method, color coded by the filtering method that retained it. Dots with grey edge, black edge were flagged by SADE as cellular-debris, invalid entropy respectively. **Right panel**: venn diagram showing agreement and disagreement among filtered 2323 barcodes between SADE, CellRanger and ArchR-TSSe. Numbers in each section are the number of barcodes in each intersection.

**Sup fig 2.**

A. Line plot of maximum read depth along chr1 of cellular-debris-containing droplet barcodes (top row), randomly selected 10 high-quality nuclei barcodes that were in agreement among all three filtering methods. Each line represents one droplet, labeled by its corresponding barcode.

B. Line plot of maximum read depth along chr1 of cellular-debris-containing droplet barcodes, with one barcode per row. Top 2 tracks (navy) were randomly selected from barcodes in agreement as reference. Bottom 3 tracks (yellow) were randomly selected from CellRanger permitted barcodes and were flagged as cellular-debris-containing droplets by SADE.

C. Fragment length distribution (FLD) for each barcode that SADE-flagged as cellular-debris-containing droplet and was deemed as high-quality by CellRanger.

**Sup fig 3.**

A. Line plot of maximum read depth along chr1 for barcodes with invalid entropy values. The top row shows barcodes uniquely permitted by ArchR-TSSe; the second row shows barcodes uniquely permitted by CellRanger; and the third row shows barcodes permitted by both methods. Each line represents one droplet and is labeled by its barcode. The bottom row serves as a reference and shows 10 randomly selected barcodes with valid entropy values that were accepted by all three filtering methods.

B. Line plot of maximum read depth along chr1 for invalid-entropy barcodes, with one droplet per row. The top two tracks (navy) were randomly selected from barcodes agreed upon by all methods and used as references. The bottom three tracks were randomly selected from barcodes uniquely permitted by CellRanger or ArchR-TSSe.

C. Fragment length distribution (FLD) for each barcode that SADE-flagged as invalid-entropy droplet and was deemed as high-quality by CellRanger or ArchR-TSSe.

**Sup fig 4.**

A. Fragment-length distributions for reads from the corresponding barcode sets, with barcodes color-coded by the filtering method that retained them.

B. Violin plot of peak-size distributions across peak sets, color-coded by the barcode set used for peak inference.

C. Scatter plot of FRIP score differences for the 2384 barcodes. The x-axis shows the difference between FRIP scores computed using SADE peaks and raw peaks, and the y-axis shows the difference between FRIP scores computed using ArchR-TSSe peaks and raw peaks. The diagonal line $y = x$ is shown for reference. Each dot represents a barcode and is colored by the filtering method that retained it: blue, uniquely retained by SADE; yellow, uniquely retained by Cell Ranger; green, retained by either Cell Ranger or SADE; and pink, uniquely retained by ArchR-TSSe.

D. FRIP scores for each peak set in barcodes identified as high-quality nuclei by at least one filtering method. Each line represents one barcode and is colored by the method that retained it, using the same color scheme as in panel C.

E. Bar plot showing the frequency of cCREs that have overlapped with peaks, with cCREs colored by class. Blue denotes peaks with no registered cCRE overlap; each such peak is counted once as "Missing cCREs." Because a peak can overlap multiple cCREs, all overlapping cCREs are counted.

F. Scatter plot of mean peak size versus the mean number of overlapping cCREs per peak for each peak set or subset. Light blue indicates raw peaks; yellow, blue, and pink indicate newly discovered peaks from Cell Ranger, SADE, and ArchR-TSSe filtering, respectively.

**Sup fig 5.** UMAP low-dimensional representation of the RNA modality gene expression of the 2381 barcodes colored by RNA clusters, or marker genes respectively.

**Sup fig 6.** A. UMAP of RNA profiles for the 2381 barcodes, colored by the ATAC filtering method that retained each barcode.

B. UMAP of RNA profiles for the 2381 barcodes, colored by its corresponding ATAC fragment read depth.

C. Scatter plot of TSS enrichment scores for barcodes uniquely retained by ArchR-TSSe. The $x$-axis shows TSSe scores calculated by ArchR-TSSe, and the $y$-axis shows TSSe scores calculated by SnapATAC.

**Sup fig 7.**
A. UMAP low-dimensional representation of the ATAC profiles of the 2384 barcodes color coded by ATAC clusters (left panel) and RNA clusters (right panel).
B. UMAP low-dimensional representation of the ATAC profiles of the 2384 barcodes colored by TSSe score calculated using SnapATAC.
C. UMAP low-dimensional representation of the ATAC profiles of the 2384 barcodes colored by entropy value.

**Sup fig 8.** For each of the four datasets, left panel is a Venn diagram showing agreement among the three filtering approaches; middle panel is a scatter plot of entropy against ATAC fragment read depth. Each dot is a barcode, colored by the filtering approaches that permitted it as quality nuclei; right panel is a scatter plot of FRIP score of each barcode calculated with respect to raw peaks inferred with all reads in the experiment.

A. Mouse brain 5k data.
B. Mouse brain cryo-preserved 5k data.
C. Human PBMC 10k data.
D. Human brain 3k data.

**Sup fig 9.** SADE-retained high-quality nuclei barcodes across a range of binned genome window sizes for each dataset. Grey dashed vertical lines indicate the window sizes (bp).

### Supplementary Figures

### Sup fig 1

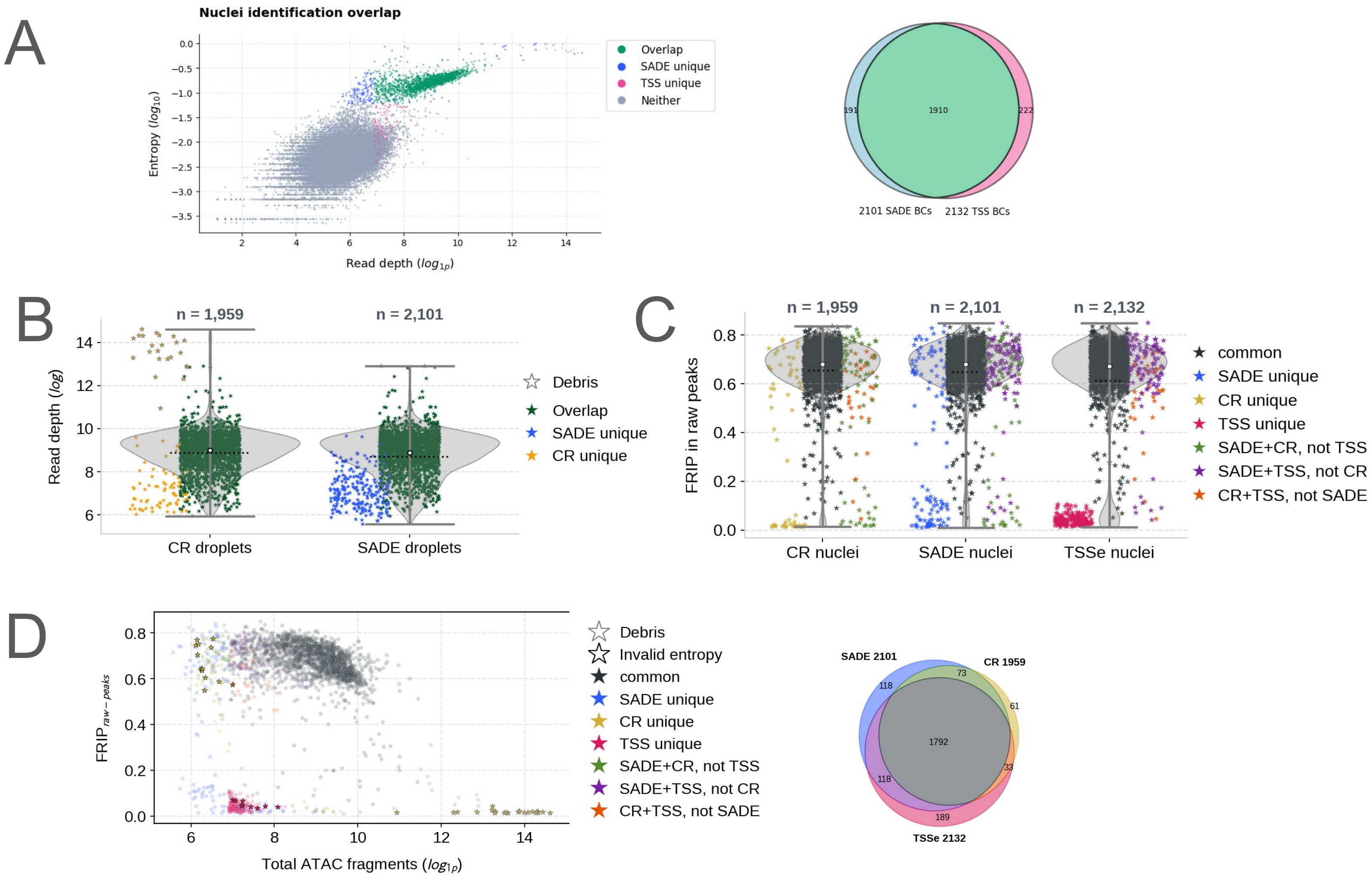
A
Nuclei identification overlap
Overlap
SADE unique
TSS unique
Neither
Entropy ($log_{10}$)
Read depth ($log_{1p}$)
191
1910
222
2101 SADE BCs
2132 TSS BCs
B
n = 1,959
n = 2,101
Read depth ($log$)
Debris
Overlap
SADE unique
CR unique
CR droplets
SADE droplets
C
n = 1,959
n = 2,101
n = 2,132
FRIP in raw peaks
common
SADE unique
CR unique
TSS unique
SADE+CR, not TSS
SADE+TSS, not CR
CR+TSS, not SADE
CR nuclei
SADE nuclei
TSSe nuclei
D
$FRIP_{raw-peaks}$
Total ATAC fragments ($log_{1p}$)
Debris
Invalid entropy
common
SADE unique
CR unique
TSS unique
SADE+CR, not TSS
SADE+TSS, not CR
CR+TSS, not SADE
SADE 2101
CR 1959
TSSe 2132
118
73
61
1792
118
33
189

### Sup fig 2

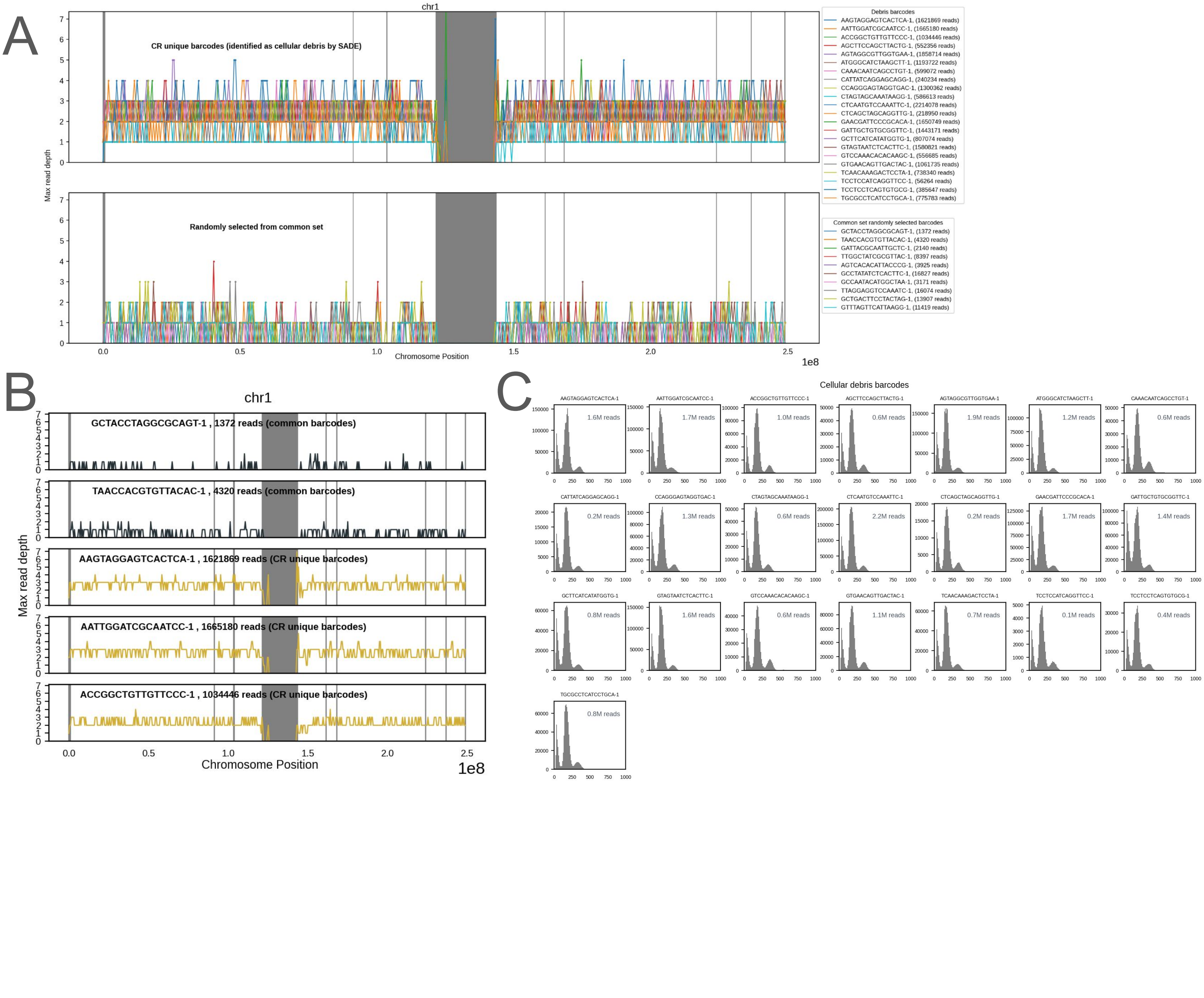
A
chr1
CR unique barcodes (identified as cellular debris by SADE)
Randomly selected from common set
Max read depth
Chromosome Position
1e8
Debris barcodes
AAGTAGGAGTCACTCA-1, (1621869 reads)
AATTGGATCGCAATCC-1, (1665180 reads)
ACCGGCTGTTGTTCCC-1, (1034446 reads)
AGCTTCCAGCTTACTG-1, (552356 reads)
AGTAGGCGTTGGTGAA-1, (1858714 reads)
ATGGGCATCTAAGCTT-1, (1193722 reads)
CAAACAATCAGCCTGT-1, (599072 reads)
CATTATCAGGAGCAGG-1, (240234 reads)
CCAGGGAGTAGGTGAC-1, (1300362 reads)
CTAGTAGCAAATAAGG-1, (586613 reads)
CTCAATGTCCAAATTC-1, (2214078 reads)
CTCAGCTAGCAGGTTG-1, (218950 reads)
GAACGATTCCCGCACA-1, (1650749 reads)
GATTGCTGTGCGGTTC-1, (1443171 reads)
GCTTCATCATATGGTG-1, (807074 reads)
GTAGTAATCTCACTTC-1, (1580821 reads)
GTCCAAACACACAAGC-1, (556685 reads)
GTGAACAGTTGACTAC-1, (1061735 reads)
TCAACAAAGACTCCTA-1, (738340 reads)
TCCTCCATCAGGTTCC-1, (56264 reads)
TCCTCCTCAGTGTGCG-1, (385647 reads)
TGCGCCTCATCCTGCA-1, (775783 reads)
Common set randomly selected barcodes
GCTACCTAGGCGCAGT-1, (1372 reads)
TAACCACGTGTTACAC-1, (4320 reads)
GATTACGCAATTGCTC-1, (2140 reads)
TTGGCTATCGCGTTAC-1, (8397 reads)
AGTCACACATTACCCG-1, (3925 reads)
GCCTATATCTCACTTC-1, (16827 reads)
GCCAATACATGGCTAA-1, (3171 reads)
TTAGGAGGTCCAAATC-1, (16074 reads)
GCTGACTTCCTACTAG-1, (13907 reads)
GTTTAGTTCATTAAGG-1, (11419 reads)
B
chr1
GCTACCTAGGCGCAGT-1 , 1372 reads (common barcodes)
TAACCACGTGTTACAC-1 , 4320 reads (common barcodes)
AAGTAGGAGTCACTCA-1 , 1621869 reads (CR unique barcodes)
AATTGGATCGCAATCC-1 , 1665180 reads (CR unique barcodes)
ACCGGCTGTTGTTCCC-1 , 1034446 reads (CR unique barcodes)
Max read depth
Chromosome Position
1e8
C
Cellular debris barcodes
1.6M reads
1.7M reads
1.0M reads
0.6M reads
1.9M reads
1.2M reads
0.6M reads
0.2M reads
1.3M reads
0.6M reads
2.2M reads
0.2M reads
1.7M reads
1.4M reads
0.8M reads
1.6M reads
0.6M reads
1.1M reads
0.7M reads
0.1M reads
0.4M reads
0.8M reads

### Sup fig 3

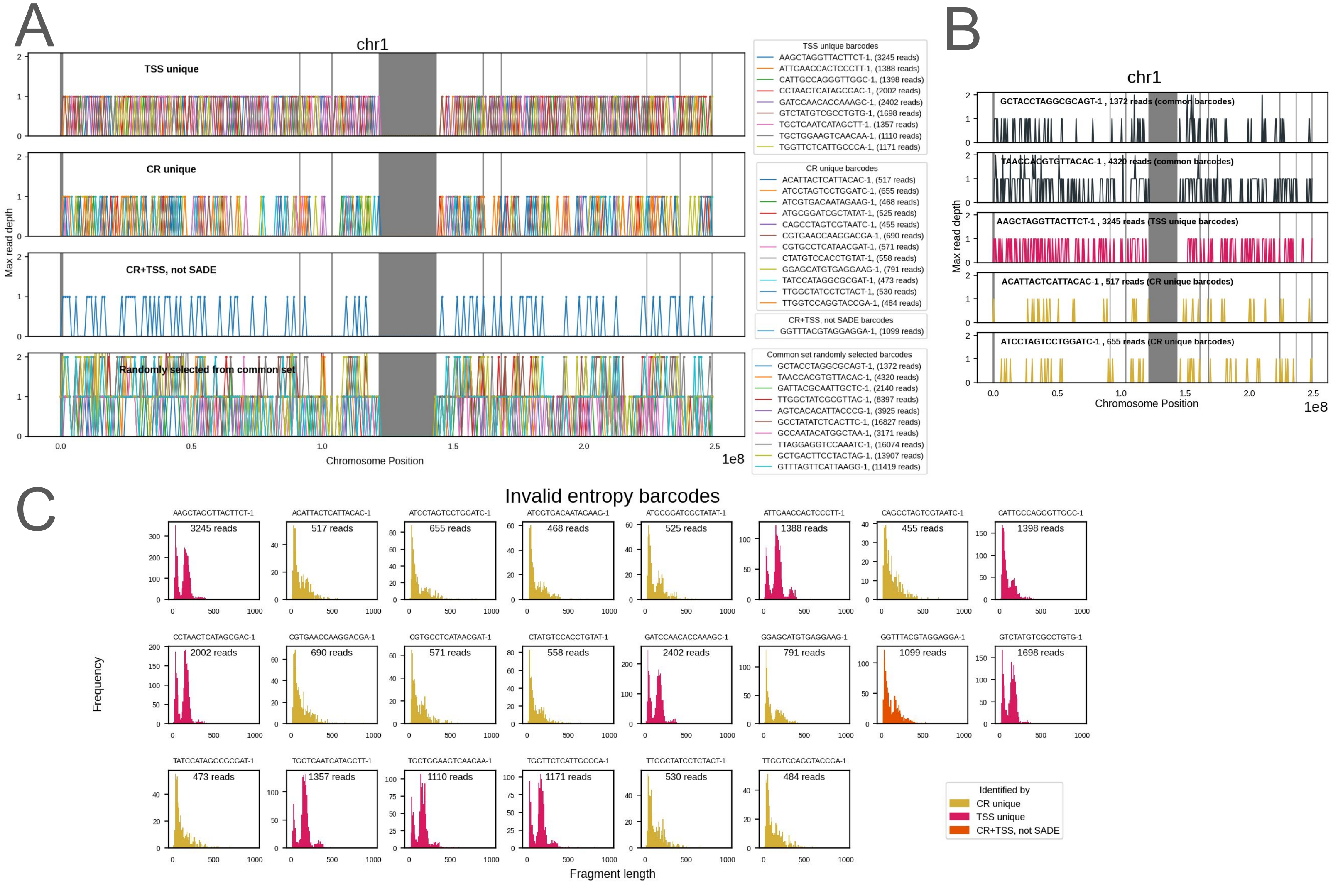
A
chr1
TSS unique
CR unique
CR+TSS, not SADE
Randomly selected from common set
Max read depth
Chromosome Position
1e8
TSS unique barcodes
AAGCTAGGTTACTTCT-1, (3245 reads)
ATTGAACCACTCCCTT-1, (1388 reads)
CATTGCCAGGGTTGGC-1, (1398 reads)
CCTAACTCATAGCGAC-1, (2002 reads)
GATCCAACACCAAAGC-1, (2402 reads)
GTCTATGTCGCCTGTG-1, (1698 reads)
TGCTCAATCATAGCTT-1, (1357 reads)
TGCTGGAAGTCAACAA-1, (1110 reads)
TGGTTCTCATTGCCCA-1, (1171 reads)
CR unique barcodes
ACATTACTCATTACAC-1, (517 reads)
ATCCTAGTCCTGGATC-1, (655 reads)
ATCGTGACAATAGAAG-1, (468 reads)
ATGCGGATCGCTATAT-1, (525 reads)
CAGCCTAGTCGTAATC-1, (455 reads)
CGTGAACCAAGGACGA-1, (690 reads)
CGTGCCTCATAACGAT-1, (571 reads)
CTATGTCCACCTGTAT-1, (558 reads)
GGAGCATGTGAGGAAG-1, (791 reads)
TATCCATAGGCGCGAT-1, (473 reads)
TTGGCTATCCTCTACT-1, (530 reads)
TTGGTCCAGGTACCGA-1, (484 reads)
CR+TSS, not SADE barcodes
GGTTTACGTAGGAGGA-1, (1099 reads)
Common set randomly selected barcodes
GCTACCTAGGCGCAGT-1, (1372 reads)
TAACCACGTGTTACAC-1, (4320 reads)
GATTACGCAATTGCTC-1, (2140 reads)
TTGGCTATCGCGTTAC-1, (8397 reads)
AGTCACACATTACCCG-1, (3925 reads)
GCCTATATCTCACTTC-1, (16827 reads)
GCCAATACATGGCTAA-1, (3171 reads)
TTAGGAGGTCCAAATC-1, (16074 reads)
GCTGACTTCCTACTAG-1, (13907 reads)
GTTTAGTTCATTAAGG-1, (11419 reads)
B
chr1
GCTACCTAGGCGCAGT-1 , 1372 reads (common barcodes)
TAACCACGTGTTACAC-1 , 4320 reads (common barcodes)
AAGCTAGGTTACTTCT-1 , 3245 reads (TSS unique barcodes)
ACATTACTCATTACAC-1 , 517 reads (CR unique barcodes)
ATCCTAGTCCTGGATC-1 , 655 reads (CR unique barcodes)
Max read depth
Chromosome Position
1e8
C
Invalid entropy barcodes
AAGCTAGGTTACTTCT-1 3245 reads
ACATTACTCATTACAC-1 517 reads
ATCCTAGTCCTGGATC-1 655 reads
ATCGTGACAATAGAAG-1 468 reads
ATGCGGATCGCTATAT-1 525 reads
ATTGAACCACTCCCTT-1 1388 reads
CAGCCTAGTCGTAATC-1 455 reads
CATTGCCAGGGTTGGC-1 1398 reads
CCTAACTCATAGCGAC-1 2002 reads
CGTGAACCAAGGACGA-1 690 reads
CGTGCCTCATAACGAT-1 571 reads
CTATGTCCACCTGTAT-1 558 reads
GATCCAACACCAAAGC-1 2402 reads
GGAGCATGTGAGGAAG-1 791 reads
GGTTTACGTAGGAGGA-1 1099 reads
GTCTATGTCGCCTGTG-1 1698 reads
TATCCATAGGCGCGAT-1 473 reads
TGCTCAATCATAGCTT-1 1357 reads
TGCTGGAAGTCAACAA-1 1110 reads
TGGTTCTCATTGCCCA-1 1171 reads
TTGGCTATCCTCTACT-1 530 reads
TTGGTCCAGGTACCGA-1 484 reads
Frequency
Fragment length
Identified by
CR unique
TSS unique
CR+TSS, not SADE

Sup fig 4

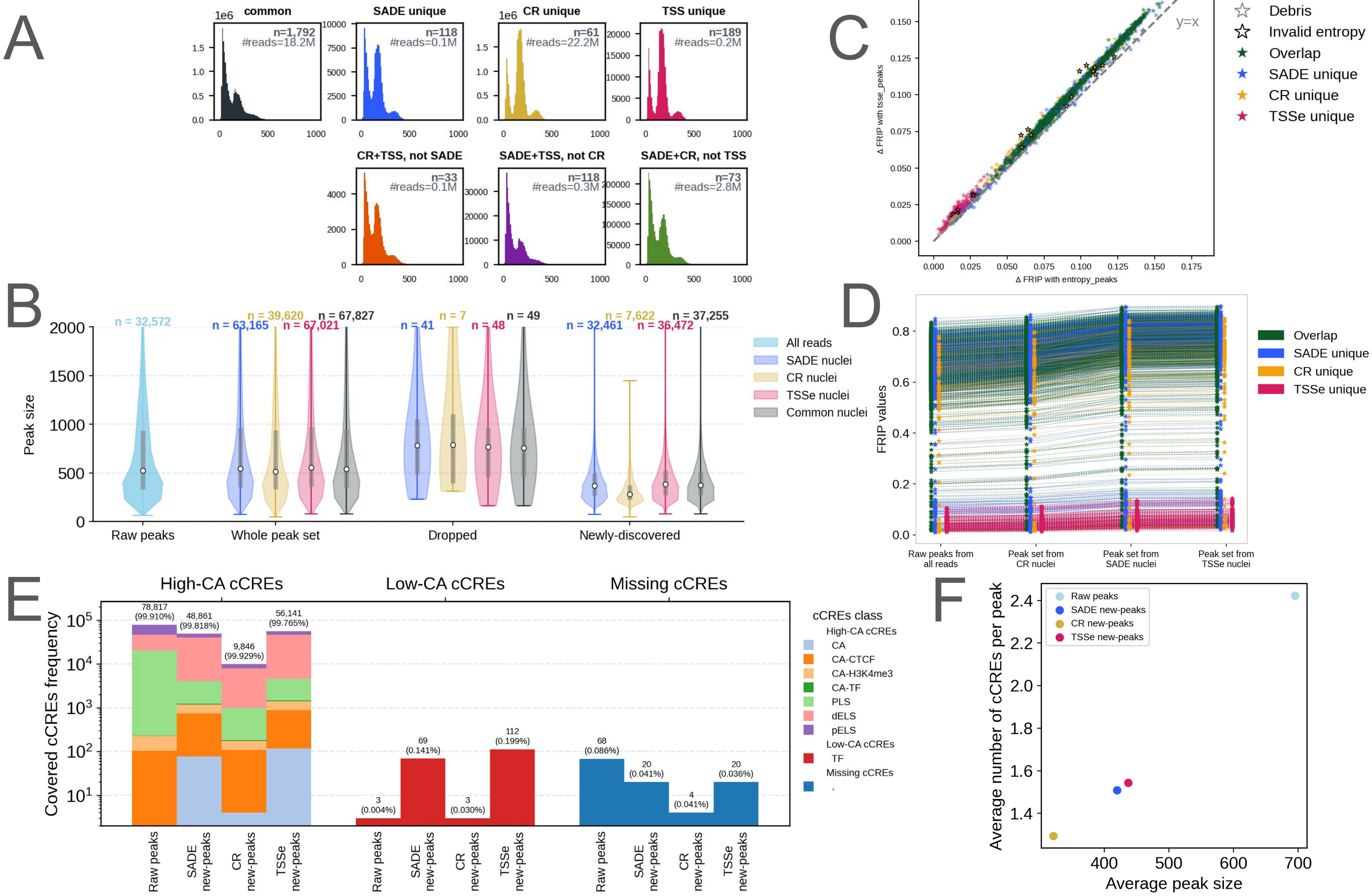

A
common
SADE unique
CR unique
TSS unique
CR+TSS, not SADE
SADE+TSS, not CR
SADE+CR, not TSS
C
Debris
Invalid entropy
Overlap
SADE unique
CR unique
TSSe unique
y=x
Δ FRIP with entropy_peaks
Δ FRIP with tsse_peaks
B
Peak size
Raw peaks
Whole peak set
Dropped
Newly-discovered
All reads
SADE nuclei
CR nuclei
TSSe nuclei
Common nuclei
D
FRIP values
Raw peaks from all reads
Peak set from CR nuclei
Peak set from SADE nuclei
Peak set from TSSe nuclei
E
High-CA cCREs
Low-CA cCREs
Missing cCREs
Covered cCREs frequency
cCREs class
F
Average number of cCREs per peak
Average peak size
Raw peaks
SADE new-peaks
CR new-peaks
TSSe new-peaks

### Sup fig 5

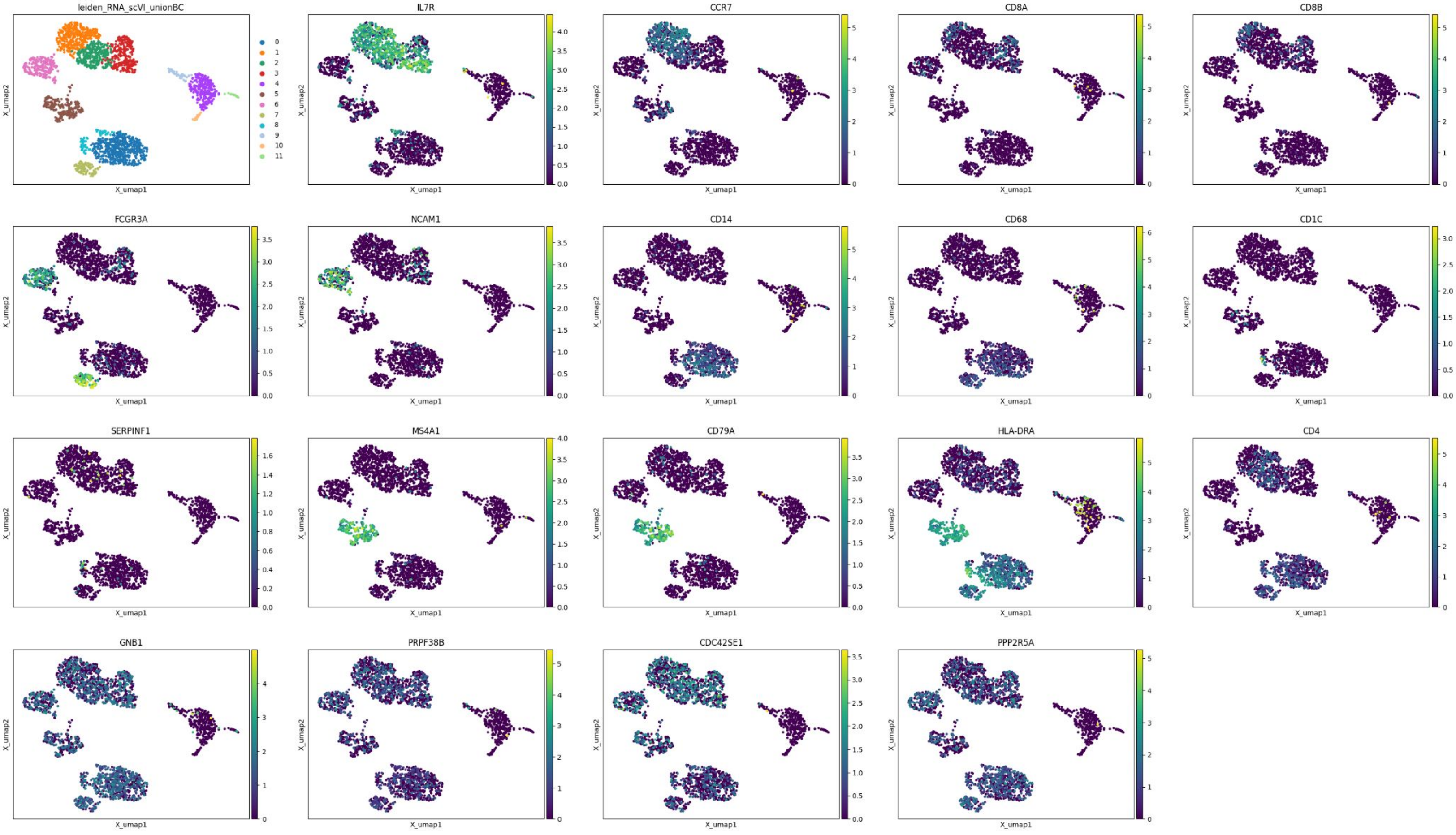
leiden_RNA_scVI_unionBC
0
1
2
3
4
5
6
7
8
9
10
11
IL7R
CCR7
CD8A
CD8B
FCGR3A
NCAM1
CD14
CD68
CD1C
SERPINF1
MS4A1
CD79A
HLA-DRA
CD4
GNB1
PRPF38B
CDC42SE1
PPP2R5A
X_umap1
X_umap2

### Sup fig 6

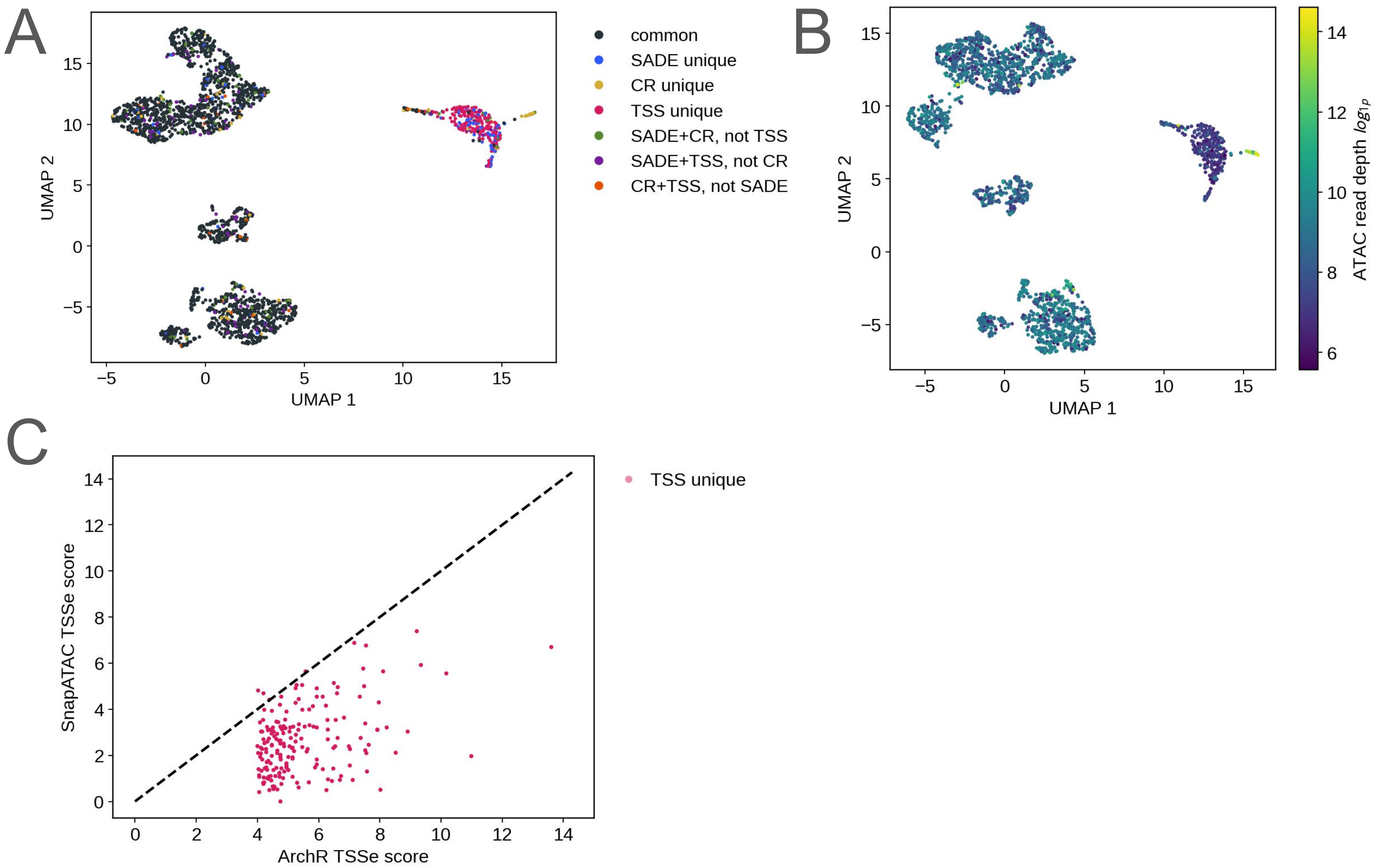
A
UMAP 1
UMAP 2
common
SADE unique
CR unique
TSS unique
SADE+CR, not TSS
SADE+TSS, not CR
CR+TSS, not SADE
B
UMAP 1
UMAP 2
ATAC read depth log1p
C
ArchR TSSe score
SnapATAC TSSe score
TSS unique

Sup fig 7

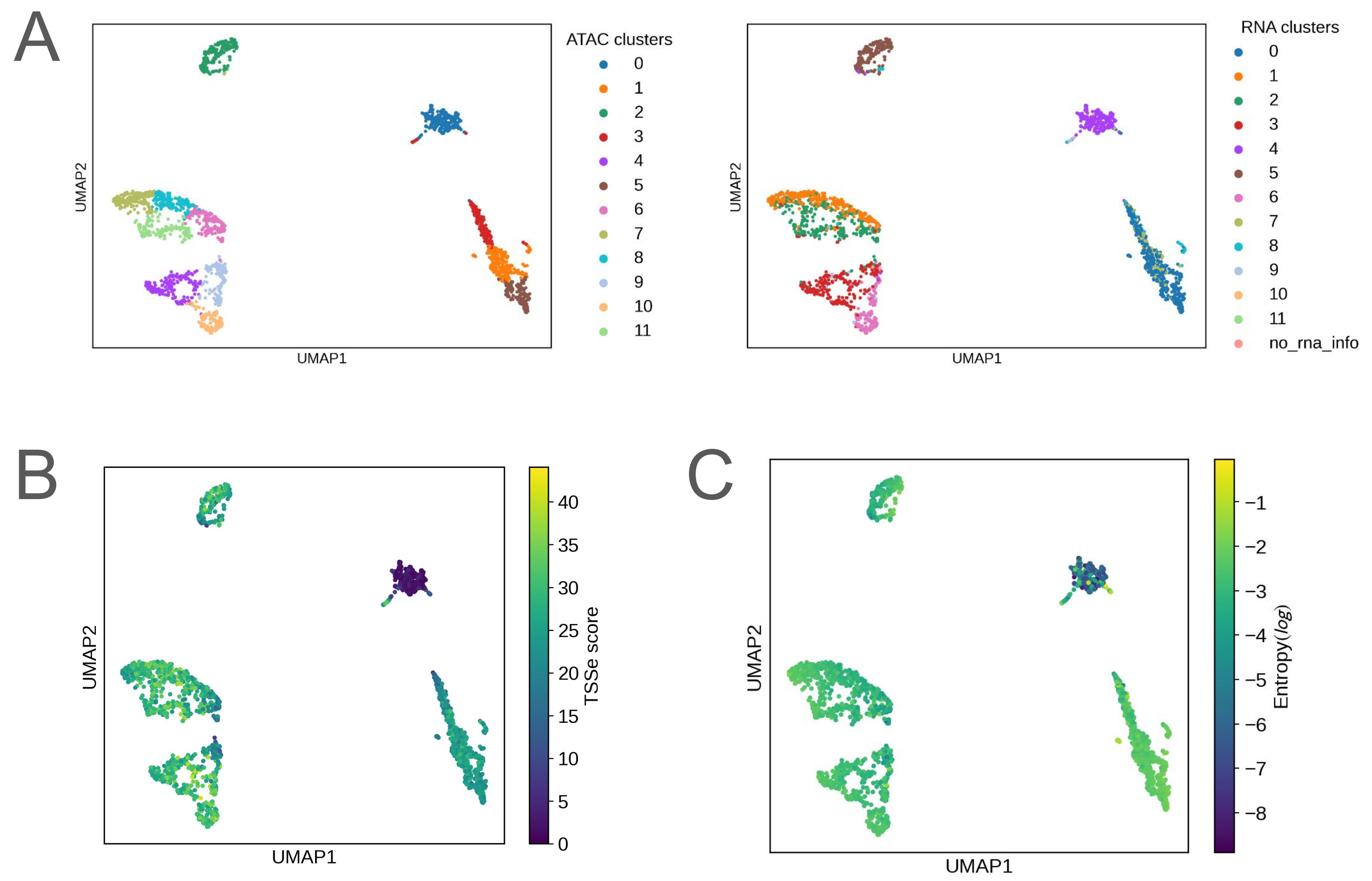
A
ATAC clusters
0
1
2
3
4
5
6
7
8
9
10
11
UMAP1
UMAP2
RNA clusters
0
1
2
3
4
5
6
7
8
9
10
11
no_rna_info
UMAP1
UMAP2
B
UMAP1
UMAP2
TSSe score
0
5
10
15
20
25
30
35
40
C
UMAP1
UMAP2
Entropy(log)
−1
−2
−3
−4
−5
−6
−7
−8

### Sup fig 8

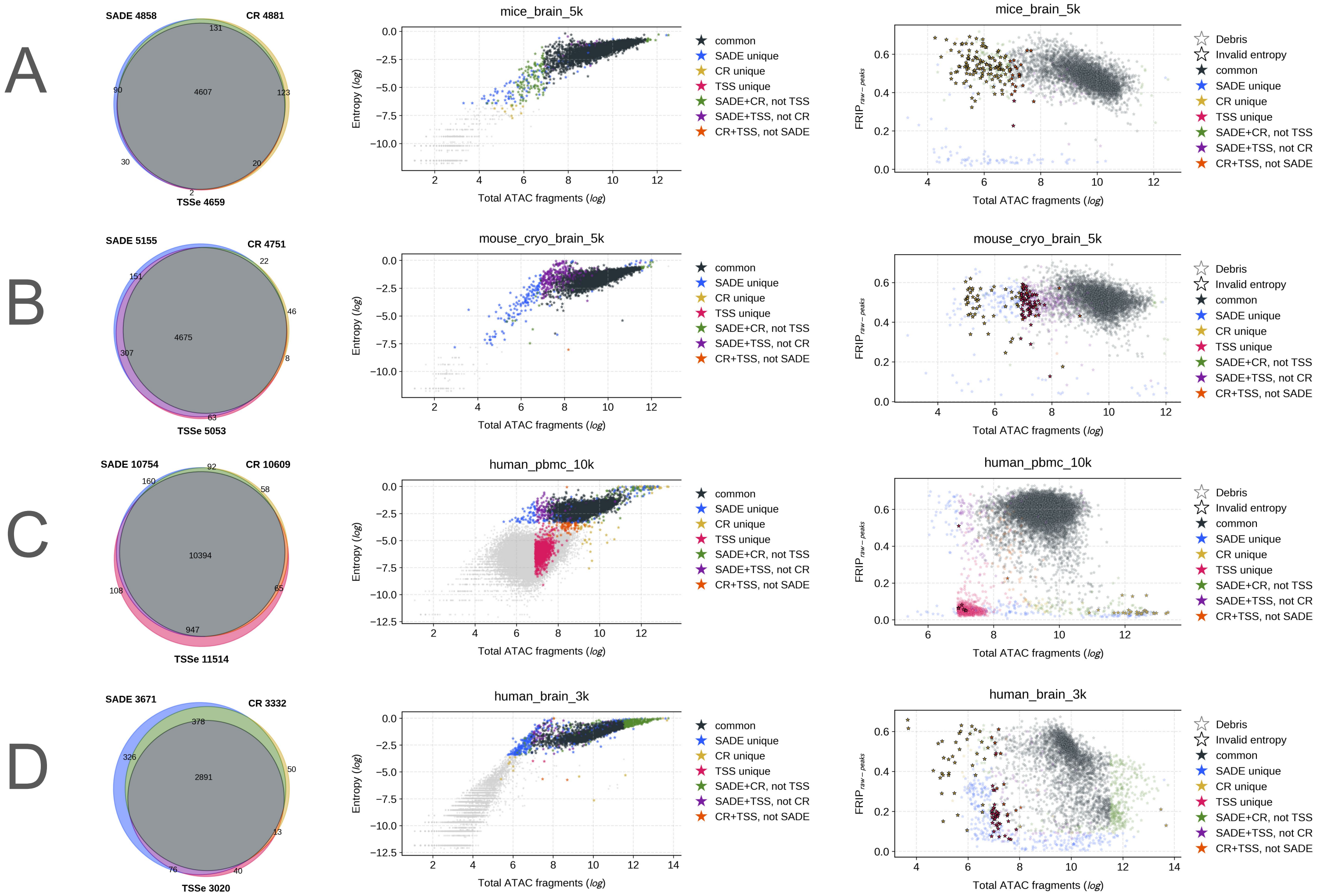

A
SADE 4858
CR 4881
TSSe 4659
4607
mice_brain_5k
B
SADE 5155
CR 4751
TSSe 5053
4675
mouse_cryo_brain_5k
C
SADE 10754
CR 10609
TSSe 11514
10394
human_pbmc_10k
D
SADE 3671
CR 3332
TSSe 3020
2891
human_brain_3k
Entropy (log)
Total ATAC fragments (log)
FRIP raw−peaks
common
SADE unique
CR unique
TSS unique
SADE+CR, not TSS
SADE+TSS, not CR
CR+TSS, not SADE
Debris
Invalid entropy

Sup fig 9

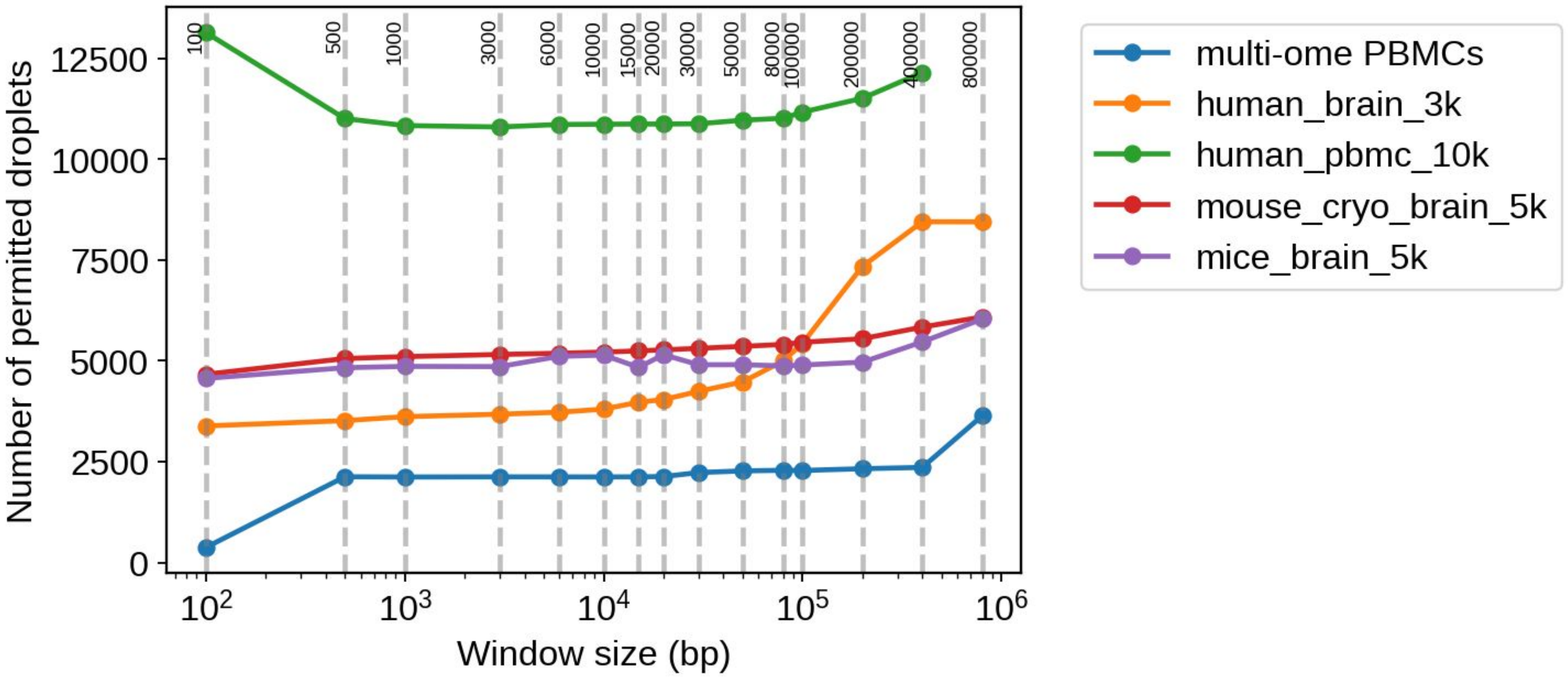

Number of permitted droplets
Window size (bp)
multi-ome PBMCs
human_brain_3k
human_pbmc_10k
mouse_cryo_brain_5k
mice_brain_5k
100
500
1000
3000
6000
10000
15000
20000
30000
50000
80000
100000
200000
400000
800000
